# Kinome-centric pharmacoproteomics identifies signaling pathways underlying cellular responses to targeted cancer drugs

**DOI:** 10.1101/849281

**Authors:** Martin Golkowski, Ho-Tak Lau, Marina Chan, Heidi Kenerson, Venkata Narayana Vidadala, Anna Shoemaker, Dustin J. Maly, Raymond S. Yeung, Taranjit S. Gujral, Shao-En Ong

**Author notes:** Correspondence (S-E.O.), (T.S.G.).

## Abstract

Kinase-dependent signaling networks are frequently dysregulated in cancer, driving disease progression. While kinase inhibition has become an important therapeutic approach many cancers resist drug treatment. Therefore, we need both reliable biomarkers that predict drug responses and new targets to overcome drug resistance. Determining the kinase(s) that control cancer progression in individual cancers can pose a significant challenge. Genomics has identified important, yet limited numbers of kinase driver mutations. Transcriptomics can quantify aberrant gene expression, but it cannot measure the protein phosphorylation that regulates kinase-dependent signaling network activity. Proteomics measures protein expression and phosphorylation and, therefore, quantifies aberrant signaling network activity directly. We developed a kinome-centric pharmacoproteomics platform to study signaling pathways that determine cancer drug response. Using hepatocellular carcinoma (HCC) as our model, we determined kinome activity with kinobead/LC-MS profiling, and screened 299 kinase inhibitors for growth inhibition. Integrating kinome activity with drug responses, we obtained a comprehensive database of predictive biomarkers, and kinase targets that promote drug sensitivity and resistance. Our dataset specified pathway-based biomarkers for the clinical HCC drugs sorafenib, regorafenib and lenvatinib, and we found these biomarkers enriched in human HCC specimens. Strikingly, our database also revealed signaling pathways that promote HCC cell epithelial-mesenchymal transition (EMT) and drug resistance, and that NUAK1 and NUAK2 regulate these pathways. Inhibition of these kinases reversed the EMT and sensitized HCC cells to kinase inhibition. These results demonstrate that our kinome pharmacoproteomics platform discovers both predictive biomarkers for personalized oncology and novel cancer drug targets.

## INTRODUCTION

Precision oncology seeks to identify specific vulnerabilities in a patient’s cancer to guide efficient treatments. Cancer genomic sequencing is the most widely used approach but has so far identified few driver mutations that can be targeted with FDA-approved drugs (*1*). Large pan-cancer pharmacogenomic screens in cell lines have revealed that in the absence of known genetic alterations, gene expression is a better predictor of drug sensitivity (*2–4*). The proteome and its post-translational modifications (PTMs) determine the activation state of cellular signaling pathways and biological processes and would likely provide a richer source of biomarkers and drug targets (*5*). Global, unbiased mass spectrometry (MS)-based proteomics directly measures protein expression and their PTMs (*6*), however, the cell signaling enzymes that process PTMs are typically underrepresented in such studies; this is also true for protein kinases, the enzymes that regulate protein phosphorylation (*7*). Kinase-catalyzed protein phosphorylation controls most cellular processes and protein kinases are frequently dysregulated in cancer (*8*). Because of that, and because they are highly druggable, kinases emerged as the most important oncology drug targets of the 21^st^ century (*9*). We reasoned that a kinome-centric pharmacoproteomics approach could be very powerful in discerning cellular mechanisms and molecular markers of kinase inhibitor drug response. To assess this, we developed a pharmacoproteomics platform that combines detailed kinobead/LC-MS kinome activity profiling (*10*) and high-throughput screening (HTS) of a 299-member diversity library of kinase inhibitors (KIs) in an array of hepatocellular carcinoma (HCC) cell lines.

HCC has grown to become the fourth most common cause of cancer-related death worldwide (*11*) and has many etiologies, including viral hepatitis, alcoholic cirrhosis, and nonalcoholic steatohepatitis (NASH) (*12*). Genomic approaches have yet to identify druggable driver genes in HCC, possibly due to its complex epidemiological, genetic, and epigenetic background (*2, 13, 14*). Remarkably, four of the five FDA-approved systemic treatments for advanced HCC, i.e. sorafenib (*15*), regorafenib (*16*), lenvatinib (*17*) and cabozantinib (*18*), are multi-kinase inhibitors; this highlights that kinase-dependent signaling networks are extremely important to HCC progression. Yet, clinicians choose treatments based on experience rather than validated biomarkers – a likely cause of the poor response rates (10-15%) (*19*). Additionally, even in HCCs that initially respond to treatment, resistance to drugs invariably develops; this is particularly well-documented for sorafenib and suggests that HCCs activate compensatory signaling pathways (*18, 20*). As a result, there is a pressing need for novel predictive biomarkers and drug targets, and we hypothesize that aberrant activities of kinases can serve as important bellwethers of drug response.

Consequently, we used our integrated pharmacoproteomics platform to study a panel of 17 diverse HCC cell lines. Kinobead/LC-MS analysis quantified 346 protein kinases, 886 kinase complex components and 11204 of their phosphorylation sites, thus specifying the activities of 284 of these signaling enzymes. Inhibitor screening generated dose response curves for 299 KIs, identifying drugs that act on specific cell lines, and compounds that broadly inhibit HCC cell growth. We correlated kinome features to drug responses and ranked proteomic features according to their association with drug sensitivity and resistance, therefore identifying molecular markers of drug response. More importantly, to obtain pathway-based drug response markers, we applied gene set enrichment analyses (GSEA) using pathways from the Reactome pathway database as gene sets (*21, 22*). This analysis identified signaling pathways that could explain either sensitivity or resistance to individual drugs. For instance, we discovered that growth inhibition by the clinical HCC drugs sorafenib, regorafenib and lenvatinib, and many other inhibitors of mitogenic and cell cycle kinases, correlated with FGFR, RAF-MEK-ERK and cell cycle pathway activation. In stark contrast, resistance to these drugs correlated with signaling pathways involved in the epithelial-mesenchymal transition (EMT) (*23*). Because kinases that regulate EMT pathways can be targeted to break drug resistance, we studied the role of EMT in HCC drug resistance in greater detail (*24*). Hence, knockdown of resistance-associated kinases revealed a FZD2-AXL-NUAK1/2 signaling module that drives HCC cell EMT and drug resistance. Pharmacological inhibition or genetic interference of AXL, NUAK1 and NUAK2 reversed the EMT, activated replication stress signaling, and sensitized HCC cells to inhibitors of checkpoint kinases (CHEK1/2) and cell cycle-dependent kinases (CDKs). Collectively, our results demonstrate that unbiased kinome pharmacoproteomics identifies molecular markers and signaling pathways that underlie drug response, reveal novel kinases important for drug resistance, and suggest rational drug combinations for the treatment of HCC. We made our dataset accessible through a web resource (*25*) that facilitates real-time interrogation and visualization.

## RESULTS

### Designing a streamlined kinome-centric pharmacoproteomics platform

Prototyping a pharmacoproteomics platform that can analyze aberrant kinome activity in an unbiased manner requires 1) that we can quantify the activation states of all identified kinases, 2) that we screen a sufficiently large and diverse set of kinase-targeted drugs to inhibit most kinases, and 3) that we use a diverse panel of cell lines that can model the activation of different kinase pathways (Figure 1A). To achieve the first, we used our kinobead/LC-MS kinome profiling platform that broadly measures kinase expression, their phosphorylation states, and interacting regulatory proteins; together, these proteomics features specify kinase activation states (Figure S1A) (*10*). To achieve the second, we used a 299-member diversity library of KIs, including selective compounds and molecules with known polypharmacology that together inhibit at least 145 primary kinase targets (Figure 1A, **Table S1**). In addition to KIs widely used as research tools, the library contained FDA-approved drugs and preclinical (Phase I-III) KIs, including the HCC drugs sorafenib, regorafenib, lenvatinib and cabozantinib. To achieve the third, we analyzed the 28 HCC cell lines from the Cancer Cell Line Encyclopedia (CCLE) (*3*) and chose 17 representative lines with diverse kinase mRNA expression (**Table S1**).

**Figure 1.**
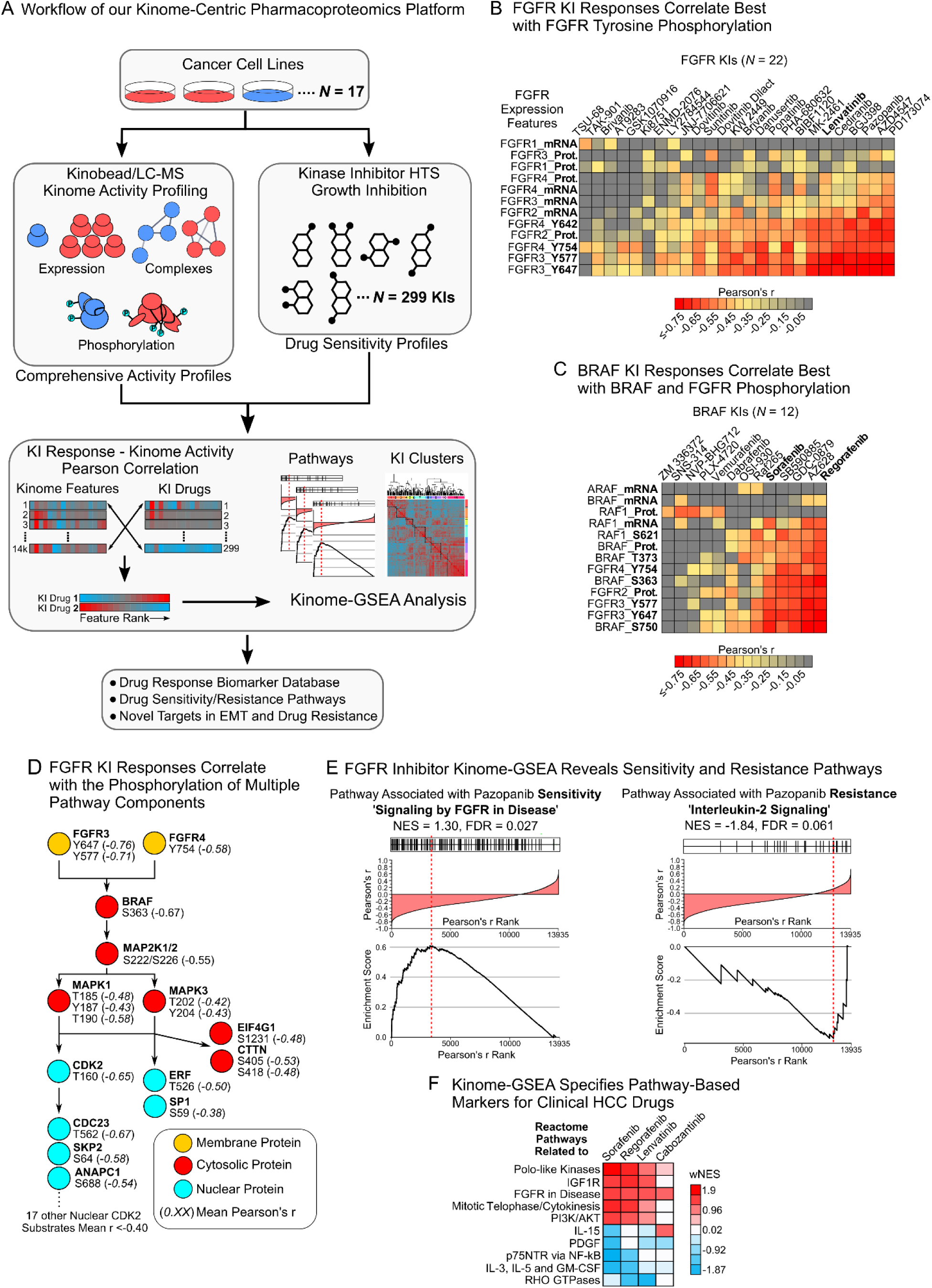
A pharmacoproteomics platform linking kinome features to drug response. (A) Schematic of our kinase-centric pharmacoproteomics platform. (B) Pearson’s r-values for FGFR1-4 expression features (mRNA, protein and phosphopeptide) correlated with the AUCs of all 22 FGFR KIs contained in our drug screen (see ‘Materials and Methods’). (C) As in (A) but for AUCs of all 12 BRAF KIs in our drug screen and RAF/FGFR expression features. (D) Phosphorylation sites in the FGFR-RAF/MEK/ERK-cell cycle signaling cascade correlated with responses to the seven FGFR KIs that showed the strongest correlation with FGFR3 and FGFR4 tyrosine phosphorylation (see (B)). Mean r-values for proteomics-AUC correlation in parentheses. (E) Kinome-GSEA identifies Reactome pathways associated with HCC cell sensitivity (left) and resistance (right) to the FGFR inhibitor pazopanib (see Figure S2D for an EGFR inhibitor example). (F) Examples for pathway that are associated with sensitivity (positive wNES) or resistance (negative wNES) to clinical HCC drugs. wNES is the FDR-weighted normalized Reactome pathway enrichment score (see ‘Materials and Methods’). See also Figure S1-3 and **Tables S1-4**.

Consequently, we used kinobead/LC-MS and label-free quantification (LFQ) to profile the kinome of the 17 HCC cell lines, and quantified 2731 proteins and 11204 phosphorylation sites. These included 346 kinases, 2821 kinase phosphorylation sites, and 886 proteins that are known to form complexes with kinases (Figure S1B and C, **Table S2**) (*26*). Functionally characterized phosphosites specified the activation states of 193 kinases (*27*) and by incorporating data on kinase complexes and their phosphorylation, we determined the activity for 284 of the 346 kinases. For instance, we detected activation of the important oncogenic kinases BRAF (e.g. S729), AKT2 (S474), the insulin-like growth factor 1 receptor (IGF1R, e.g. activating Y1164), and AXL (GAS6 complex), among many others (**Table S2**). The broad coverage of kinases demonstrates that kinobead/LC-MS is a powerful tool that readily identifies molecular mechanisms of cell signaling.

Next, we conducted a CellTiter-Glo HTS with the 299-member KI library and generated seven-point dose-response curves for each KI in all 17 HCC cell lines (**Table S3**). Using the area under the dose-response curve (AUC) as a measure of drug efficacy (low AUC – high efficacy), we observed diverse responses to inhibitors. Hence, some KIs classes, for instance FGFR, EGFR, IGF1R and BRAF inhibitors, blocked cell growth in certain HCC lines (Figure S2A). In contrast, inhibitors of MEK, cell cycle-related kinases and MTOR were broadly active, efficiently inhibiting cell growth in 10 of the 17 cell lines (Figure S2A, **Table S3**). The diversity of drug responses shows that our KI screen contained enough drugs with orthogonal target profiles to inhibit many different mitogenic and survival kinases, an important requirement for our unbiased correlation analyses.

### Correlation of kinome features with drug responses identifies predictive biomarkers

To test if our kinome-centric pharmacoproteomics platform can identify molecular markers of KI drug response, we used Pearson’s r to correlate the MS intensities of all 13935 proteomics features in our dataset (**Table S2**) with each KI’s AUC values (**Table S3**) across the 17 HCC cell line panel (*N* = 17, for details see ‘Materials and Methods’). The resulting matrix of Pearson’s r-values (13935 X 299 matrix in **Table S4**) ranks kinome features according to their correlation with drug sensitivity (negative r: high MS intensity – low AUC), or alternatively, correlation with cell survival and KI resistance (positive r: high MS intensity – high AUC); analyzing this matrix should, therefore, readily identify predictive biomarkers. Reasoning that kinase activation predicts the efficacy of inhibitors targeting that kinase, we tested the validity of our approach by examining biomarkers of the clinical HCC drug lenvatinib and 21 other FGFR inhibitors in our KI panel (Figure 1B). Gratifyingly, the activating FGFR3 phosphorylation sites Y647 and Y577, and the activating Y754 on FGFR4 correlated very well with responses to most FGFR inhibitors. In contrast, kinase protein and mRNA expression correlated less well with drug responses, as these features do not necessarily determine the kinase activation state (Figure 1B). Encouraged by these results, we examined if the HCC drugs sorafenib and regorafenib and 10 other BRAF inhibitors in our KI panel also correlated with target kinase phosphorylation (Figure 1C). Indeed, BRAF phosphorylation on S750 and S363 predicted drug response better than BRAF protein or mRNA expression. We obtained additional evidence for this high predictive power of activating kinase phosphosites by testing 26 EGFR inhibitors (Figure S2B). These drugs correlated best with activating tyrosine phosphorylation of the EGFR on Y1110 and Y1172. Collectively, these results demonstrate, first, that in many instances the phospho-activation state of kinases predicts drug response better than protein and mRNA expression and, second, that our kinome-centric pharmacoproteomics approach readily detects these predictive markers.

While analyzing BRAF inhibitor response markers, we noticed that not only BRAF phosphorylation, but also FGFR tyrosine phosphorylation correlated tightly with the response to these drugs (Figure 1C). Because BRAF is a key component of the FGFR signaling pathway, we reasoned that multiple phosphorylation events along the same signaling axis can act as drug response markers. Consequently, we analyzed the correlation of phosphosites on FGFR pathway components with responses to lenvatinib and six other FGFR inhibitors that correlated with the activation of the FGFR3 and FGFR4 (Figure 1B and D). Strikingly, this revealed that activating phosphosites on MEK1/2 and ERK1/2, kinases situated downstream of FGFR-BRAF signaling, and activating sites on the cell cycle driver CDK2 (T160) all correlated very well with FGFR inhibitor response (Figure 1D). We expanded this analysis further to include phosphosites on kinase substrates that regulate the cell cycle (e.g. CDC23-T562 and SKP2-S64), transcription (ERF-T526 and SP1-S59) and translation (EIF4G1-S1231), and indeed, these substrates also correlated with FGFR inhibitor response (Figure 1D). We obtained similar results when analyzing the correlation of EGFR pathway members with EGFR inhibitors (Figure S2C). Together, these results indicate that our kinome pharmacoproteomics approach identifies not only individual phospho-markers of drug response but can also be used to construct valuable pathway-based biomarkers.

### GSEA analysis of kinome features identifies pathway-based drug response markers

Compared to single-feature markers, pathway-based biomarkers predict patient outcomes with increased accuracy (*28, 29*). Unbiased analyses of signaling pathways in disease could also expose underlying disease mechanisms and facilitate target discovery. Because we found that FGFR inhibitor responses correlated with activation of multiple FGFR-BRAF/MEK/ERK-cell cycle pathway components (Figure 1D), we hypothesized that our kinome pharmacoproteomics platform can identify such pathway-based biomarkers. To give our pharmacoproteomics platform a quantitative and statistical framework for pathway-based biomarker discovery, we applied an unbiased GSEA analysis using Reactome pathways as the gene sets (*21, 29*). Importantly, because GSEA can detect pathway enrichment at the opposing ends of ranked feature lists, it can both identify pathways that correlate with drug sensitivity and drug resistance.

Consequently, we collated our kinome features into 327 cancer-relevant Reactome signaling pathways (**Table S3**) (*21*) and used a modified GSEA (*22*), hereafter referred to as kinome-GSEA, to identify pathways associated with each drug. As the result, we obtained scores for 275 of the 327 pathways, with adjusted p-value and normalized enrichment score (NES), thus ranking pathways for their association with sensitivity (positive NES) or resistance (negative NES) to each of the 299 KI drugs. To test the performance of our analysis, we first looked at pathways enriched by well-characterized KIs. For example, the selective FGFR inhibitor pazopanib enriched pathways terms associated with FGFR and cell cycle activation (Figure 1E and **Table S3**). In contrast, the selective EGFR inhibitor lapatinib enriched Reactome pathways associated with EGFR, PI3K and NF-κB activation (Figure S2D and **Table S3**). Remarkably, both inhibitors de-enriched pathways that are known to regulate tyrosine kinase inhibitor resistance in various cancers. Thus, pazopanib de-enriched pathways associated with interleukin signaling (JAK-STAT pathway, Figure 1E) (*30*), and lapatinib de-enriched Wnt-signaling pathway terms (Figure S2D) (*31*). Expanding our analysis to include the clinical HCC drugs sorafenib, regorafenib and lenvatinib (BRAF and FGFR KIs) confirmed that these drugs are effective when FGFR and cell cycle pathways are activated, and ineffective when survival pathways such as interleukin and NF-kB signaling are activated (Figure 1F). In contrast, the clinical AXL and c-Met inhibitor cabozantinib correlated less well with these pathways. Collectively, these results demonstrate that our kinome-GSEA analysis identifies signaling pathways and cellular mechanisms accounting for kinase inhibitor sensitivity and resistance. These pathways may serve as predictive biomarkers in the clinic and may specify novel targets for HCC treatment.

### Kinome-GSEA outperforms mRNA-based analyses in predicting drug target pathways

Our kinome-GSEA analyses uses proteomics features of kinase activation, including protein expression and phosphorylation, to identify pathway-based markers of drug response. Naturally, the question arose how this approach performs when compared to similar analyses with transcriptomic data. Therefore, we ran the GSEA using CCLE mRNA data of the 17 HCC cell lines as the input (Figures 2A and S3, see also ‘Materials and Methods’) (*3*). To compare performance between kinome- and mRNA-GSEA directly, we extracted the FDR-weighted normalized pathway enrichment scores (wNES) for a set of 11 well-defined KI classes (Figure 2A). We assumed that Reactome pathway gene sets containing the primary KI target were the most indicative of drug response and compared wNES values for these pathways (Figure 2A). This revealed that most KI classes (9 of 11) were best described by pathways enriched by kinome-GSEA analysis, while mRNA data yielded better results than the kinobeads for the remaining 2 KI classes. For instance, kinome-GSEA accurately identified drug sensitivity pathways containing the IGF1R or EGFR gene name for selective inhibitors of these kinases, while the corresponding mRNA-GSEA performed poorly (Figure 2A). In contrast, mRNA-GSEA was better at identifying the signaling pathways relevant to AURKA and MTOR inhibitor response. Our results suggest that kinobead/LC-MS analysis would outperform mRNA profiling in detecting pathway-based biomarkers for most KI drugs.

**Figure 2.**
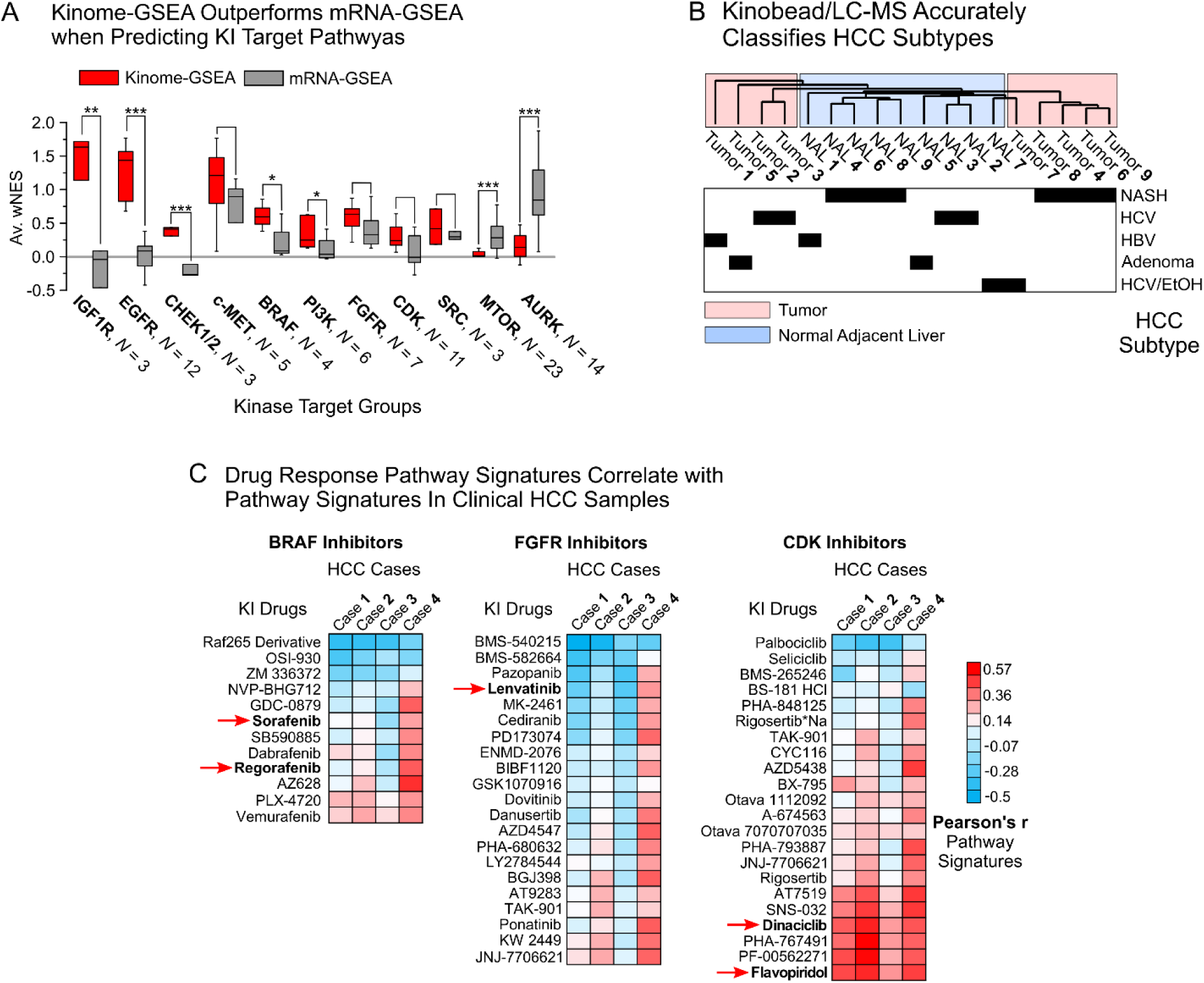
Kinome-GSEA performance and kinobead/LC-MS profiling of clinical HCC specimens. (A) GSEA analysis using kinobead/LC-MS data compared to CCLE mRNA data to identify signaling pathways that are the targets of selected KI groups. Kinome-GSEA analysis outperforms mRNA analysis for 9 of 11 important KI groups. For an analysis of all KIs see Figure S3 (see also ‘Materials and Methods’). *: p < 0.1, **: p < 0.01, ***: p < 0.001 from a two-sample Student’s T-test. (B) Hierarchical clustering of the 9 paired tumor/NAL HCC samples by kinome expression features as determined by super-SILAC kinobead/LC-MS profiling (see also ‘Materials and Methods’). (C) We correlated KI drug response marker signatures specified in 17 HCC lines with kinome pathway signatures identified in clinical HCC samples (kinome-GSEA wNES values), showing that pathway-based markers for specific KI drugs are enriched in human tumors. BRAF, FGFR and CDK target groups are shown. Highlighted drugs are used to treat HCC in the clinic of undergo clinical trials (see **Table S5** and ‘Materials and Methods’). See also Figure S3, and **Table 5**.

**Figure 3.**
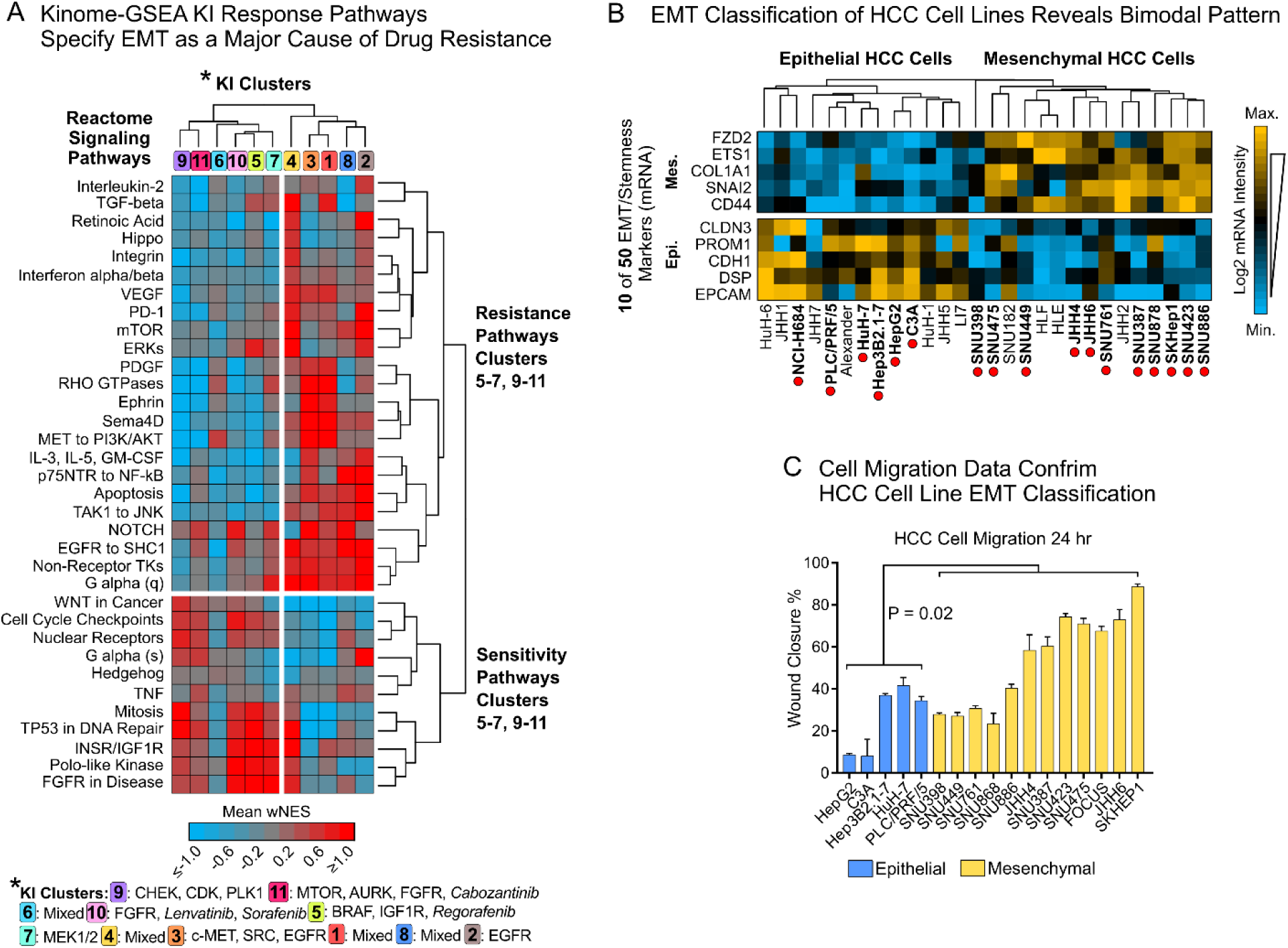
The mesenchymal-epithelial transition (EMT) state specifies HCC resistance to kinase inhibition. (A) Heatmap of average wNES values for the KI drug clusters identified by Pearson correlation and clustering of Reactome pathway wNES values for the 299 KIs (see **Figure S4D** and **Table S3**). 34 representative pathways with the highest range in average wNES across the 11 KI clusters are shown (see ‘Materials and Methods’). Abbreviated Reactome pathway term shown (for a complete list of pathways see **Table S3**). (B) Hierarchical clustering of EMT marker mRNA expression in the 28 CCLE HCC cell lines. (C) Difference in cell motility between epithelial and mesenchymal HCC cells (wound healing assay, two-tailed Student’s T-test, P = 0.02). See also Figure S4, and **Table S1-4**.

### Pathway-based kinome-GSEA biomarkers are enriched in clinical HCC specimens

Having established the performance of our kinome-pharmacoproteomics platform in HCC cell lines, we next evaluated if our pathway-based biomarkers sets can be detected in clinical HCC specimens.

First, we tested if our kinobead/LC-MS kinome profiling workflow (*10*) can be applied to clinical tumor tissue samples. To that end, we profiled nine paired HCC tissue samples (tumor and normal adjacent liver, NAL) together with a super-SILAC lysate standard (see ‘Materials and Methods’) (*32*), quantifying 251 protein kinases and 1172 other proteins (**Table S5**). Remarkably, clustering the 18 tumor and NAL samples by protein expression separated tumors from NAL samples and further classified HCCs by their distinct etiologies; this was particularly evident for the four tumors with underlying non-alcoholic steatohepatitis (NASH) (Figure 2B). These results confirm, first, that kinobead/LC-MS can readily quantify kinome features in human HCC specimens and, second, that these kinobead/LC-MS features accurately describe HCC etiologies.

Encouraged by these results, we analyzed four additional tumor-NAL pairs with LFQ kinobead/LC-MS, quantifying both protein expression and phosphorylation. Gratifyingly, we found that the performance of our LFQ protocol in HCC tissue specimens is comparable to our HCC cell line experiments. We quantified 2151 kinase phosphorylation sites on 286 kinases and 680 kinase interactors from 1 mg of protein extract, obtaining phosphorylation and interaction-dependent kinase activity in human tumor specimens (**Table S5**, Figure S4A-C). We anticipate that our optimized kinobead/LC-MS workflow (*10*) will enable routine kinome profiling of clinical cancer samples. Next, we applied kinome-GSEA analysis to identify Reactome pathways that are upregulated in tumors or NAL samples (**Table S5**). To test if pathway-based drug response markers identified from cell lines are enriched in clinical HCC samples, we correlated the pathway signatures, i.e. the list of Reactome wNES values, of each of the 299 KIs with the signatures of each of the four clinical HCC specimens (see ‘Materials and Methods’). Strikingly, this revealed that pathway-based markers of clinical HCC drugs are highly enriched in specific tumors (Figure 2C). For instance, HCC case 4 showed high enrichment of pathways that specify sorafenib and regorafenib sensitivity (r ∼0.4), as well as sensitivity of FGFR inhibitors. Therefore, in a clinical setting, these findings would suggest that patient 4 is most likely to respond to sorafenib or regorafenib treatment. In contrast, CDK inhibitor pathway markers were highly enriched in 3 out of 4 tumors (r ∼0.4 to ∼0.5, Figure 2C), indicating that most patients would respond to CDK inhibitor treatment, for instance with dinaciclib or flavopiridol, which are currently in clinical trials for HCC (*33*). Collectively, these data establish for the first time that kinome activity-based biomarkers can be detected in human tissue specimens, possibly paving the way for future clinical trials of personalized therapies based on such pathway-based biomarkers.

### The cellular EMT state determines response to kinase inhibitors

Having established our ability to identify pathway-based biomarkers for specific HCC drugs, we next wished to explore similarities in drug response pathways among all 299 kinase inhibitors; this would identify pathways and mechanisms that regulate the response to a broad range of clinical and pre-clinical KI drugs. To detect such similarities, we used a pairwise-correlation of pathway wNES values for each KI and clustered the resultant r-values. This analysis revealed 11 clusters of KI drugs grouped by their similarities in GSEA-enriched signaling pathways (Figure S4D, **Table S3**, see also ‘Materials and Methods’). Groups of well characterized KIs are thus found among other less well characterized drugs, offering testable hypotheses about the similarities of drug action for these KI groups. For instance, the clinical HCC drugs sorafenib and lenvatinib clustered with various FGFR inhibitors (Cluster 10) and were distinct from the compounds in cluster 5, that included regorafenib, and various BRAF and IGF1R inhibitors (Figure S4D).

Next, to identify the principal kinase signaling pathways governing the response to these 11 KI drug clusters, we extracted 34 representative cancer-relevant Reactome pathways from the larger panel of 275 scored pathways (**Table S3**, see also ‘Materials and Methods’). Strikingly, clustering of kinome-GSEA enrichment scores of these 34 pathways produced a clear separation into two distinct groups (Figure 3A). KIs in clusters 5-7 and 9-11 formed one group with positive enrichment scores for pathways commonly overexpressed in rapidly proliferating cells including FGFR-, IGF1R-, cell cycle-, and mitosis-related pathways. KIs in this group mainly inhibit BRAF, the FGFR isoforms, including sorafenib, regorafenib and lenvatinib, the IGF1R, and cell cycle-related kinases (PLK1, CDKs, CHEK1/2). Concomitantly, this same group had negative enrichment scores for kinase signaling pathways related to c-MET, TGF-β, cytokine and NF-kB signaling (Figure 3A); these pathways are known to regulate the EMT, an important mechanism of drug resistance in cancer (*24, 34*). Conversely, compounds in clusters 1-4 and 8 (e.g. EGFR, c-MET and SRC KIs) showed positive enrichment scores for these same EMT-associated pathway terms and negative scores for FGFR- and cell cycle-related terms (Figure 3A). This opposing behavior of EMT pathway activation and KI drug response suggested *i*) the presence of mesenchymal-like HCC cells in our panel and *ii*) that EMT is the cause of drug resistance against a broad range of KIs.

To test if our panel contains cell lines in different EMT states, we examined the mRNA expression of 50 important EMT and stem cell markers in the 28 CCLE HCC cell lines (*3*) (**Table S1**). We found that their expression varied considerably, with 23/50 mRNAs changing >100-fold across the panel. Indeed, semi-supervised hierarchical clustering of EMT markers classified the panel into 14 epithelial and 14 mesenchymal lines (Figure 3B and **Table S1**). As EMT is typically associated with increased cell motility (*35*), we assayed the cell migration of 15 HCC lines, confirming that the mesenchymal HCC cell lines exhibited significantly enhanced wound closure compared to epithelial lines at 24 h (P = 0.02, Figure 3C). The wound healing assay data confirmed our classification of HCC cell lines into distinct epithelial and mesenchymal groups and provides further evidence that the EMT is a major cause of HCC drug resistance to a broad range of KI drugs (see also Figure S2A).

### A comprehensive map of kinases associated with HCC cell EMT state and drug response

An important next step in our analysis was to identify the kinases and pathways associated with the HCC cell EMT state and drug response phenotypes; such kinases could represent prognostic biomarkers as well as important clinical targets to reverse EMT and overcome drug resistance. Accordingly, to quantify and prioritize EMT state-specific kinome features, we applied T-test statistics to our dataset of LFQ-MS intensity values in epithelial (n = 7) vs. mesenchymal (n = 10) HCC cells (Figure 3B and **Table S2**). This analysis identified 101 kinases, 380 kinase phosphosites, and 938 other phosphoproteins that differed significantly in expression between the two EMT phenotypes (Figure 4A and **Table S2**).

**Figure 4.**
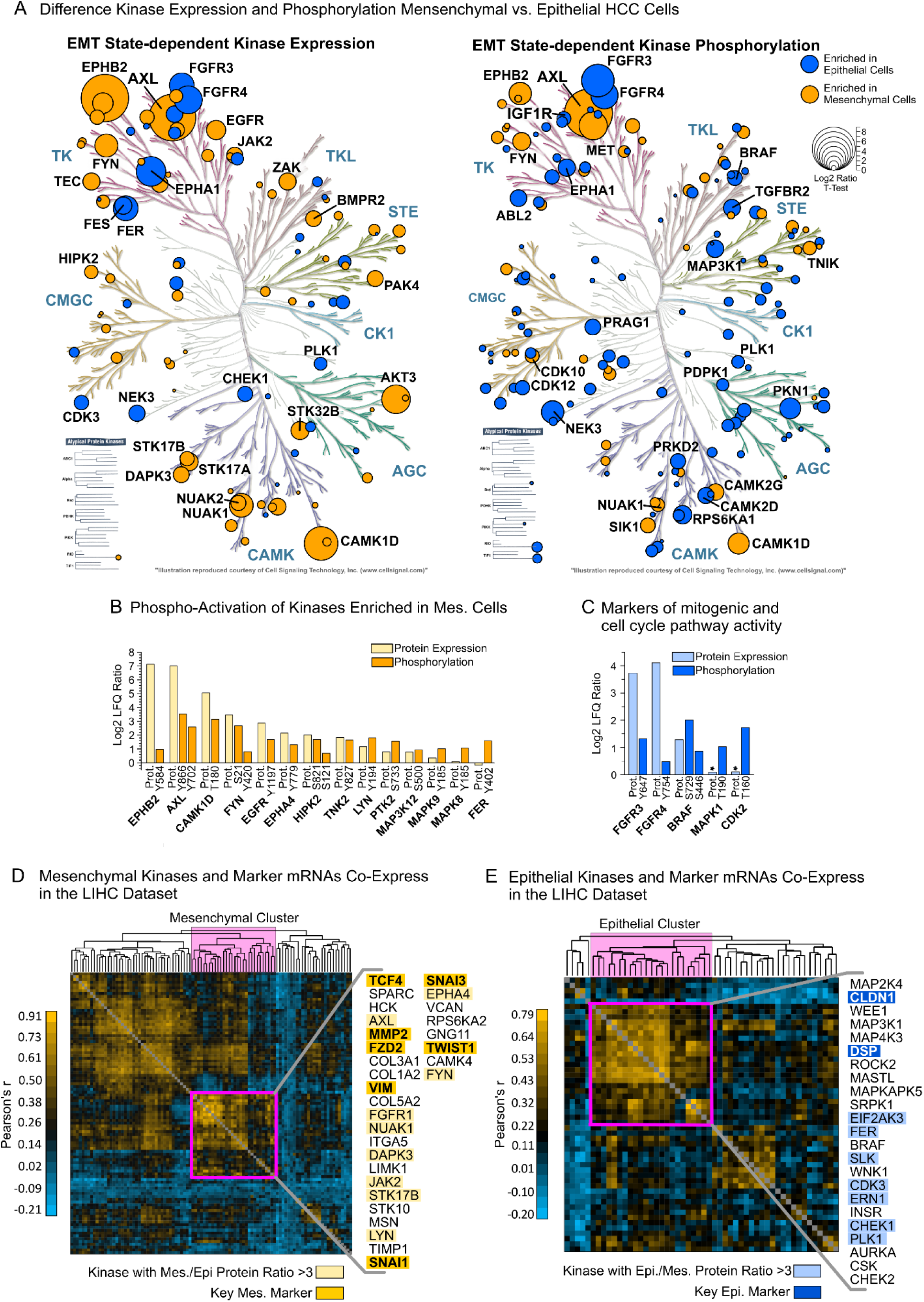
The EMT-associated kinome in HCC cells lines and Clinical HCC specimens. (A) Kinases (circles, left) and their phosphorylation sites (right) significantly overexpressed in epithelial or mesenchymal HCC cell lines (FDR = 0.05). Kinases with a log2 ratio >2 are labeled. The largest phosphosite ratio between EMT phenotypes for each kinase is plotted. (B) Activating phosphosites on kinases overexpressed in mesenchymal compared to epithelial cells. (C) Activating phosphosites on important mitogenic and cell cycle kinases overexpressed in epithelial compared to mesenchymal cells. (D) Co-expression and clustering analysis of TCGA-LIHC mRNA data of 66 mesenchymal kinases identified in the 17 HCC cell line panel and 34 mesenchymal markers (*N* = 424 samples, see ‘Materials and Methods’) (E) As in (D) but 35 epithelial kinases and 17 epithelial markers. See also **Table S2**.

Remarkably, the protein expression of 66 out of 101 EMT kinases was significantly upregulated in mesenchymal cells, including the clinically important cabozantinib targets AXL (>100-fold) and c-MET (5-fold) (*18*). Other highly regulated kinases were the RTK EPHB2 (>100-fold) and non-receptor kinases FYN, AKT3, CAMK1D, NUAK1 and NUAK2 (all > 4-fold, Figure 4A). The EMT-associated phosphokinome revealed that AXL’s kinase activity, but not c-MET’s, was increased in mesenchymal HCC cells, suggesting that AXL plays a more important role in EMT than c-MET. EPHB2, CAMK1D, FYN and 18 other kinases were also highly phosphorylated on activating sites in mesenchymal HCC cells, indicating that these kinases may play important roles in maintaining the mesenchymal state (Figure 4B and **Table S2**).

Conversely, epithelial HCC cells showed increased expression of 35 kinases, including the lenvatinib targets FGFR3 and 4 (∼15-fold), and the sorafenib and regorafenib target BRAF (3-fold, Figure 4A). Other highly upregulated kinases (> 4-fold) included NEK3, CHEK1, CDK3, and PLK1 that have important roles in cell cycle and DNA damage response (DDR) signaling. Additionally, we identified activating phosphosites on kinases and specific kinase substrates enriched in the epithelial cell phosphoproteome that confirmed increased activity of the cell cycle through CDK2 and mitogenic signaling through FGFR3/4, BRAF and MAPK1/3 (Figure 4C). These findings agree with previous reports stating that proliferation- and cell cycle-related pathways are specifically upregulated in epithelial cancer cells and downregulated during cancer cell EMT (*36*). Collectively, our results represent the first comprehensive proteomics dataset of kinase signaling associated with the HCC cell EMT state. Our dataset will serve as a valuable resource for cancer researchers that identifies the kinases and downstream signaling pathways associated with either a drug-resistant mesenchymal phenotype or proliferative drug-sensitive epithelial phenotype in HCC.

To gauge the clinical relevance of the EMT-associated kinases identified in HCC cell lines, we analyzed their mRNA expression in the TCGA-Liver Hepatocellular Carcinoma (TCGA-LIHC) dataset (*13*). Specifically, we analyzed the co-expression of 50 EMT markers and our 101 EMT kinases (**Table S1** and **S2**) across all 424 tissue samples in the LIHC dataset. This analysis identified multiple clusters of co-expressed EMT markers and kinases. For instance, among mesenchymal mRNAs, we identified a cluster containing *AXL*, *EPHB2*, *NUAK1, FYN* and key EMT drivers *TWIST1*, *SNAI1,* and *FZD2* (Figure 4D). We also identified a cluster of epithelial markers and kinases containing the HCC drug target *BRAF* as well as the cell cycle checkpoint kinases *CHEK1/2, WEE1, and PLK1* (Figure 4E). These results show that the functional relationship between EMT kinases and markers is conserved in human tumors, therefore suggesting that EMT transcriptional programs, including our EMT kinases, are active in human HCCs.

### AXL drives reprogramming of the EMT-associated kinome in HCC

Our kinome profiling data revealed that activating phosphorylation sites on AXL and the protein itself were among the most differentially expressed in mesenchymal vs. epithelial HCC cells (top and second most, respectively; Figure 4A, 4B, and 5A). Furthermore, AXL co-expressed with key EMT markers and other EMT kinases identified in human tumors (Figure 4D). AXL is an important player in cancer cell EMT (*37*) and the development of tumor metastasis and drug resistance in HCC (*38, 39*), as highlighted by the recent success of cabozantinib (AXL and c-MET inhibitor) as a second-line treatment for sorafenib-resistant HCCs (*18*). However, no detailed proteomics studies of AXL signaling have been published to date. This is surprising as such studies may reveal important AXL pathway components that can serve as clinical biomarkers of the EMT and molecular targets to break drug resistance.

**Figure 5.**
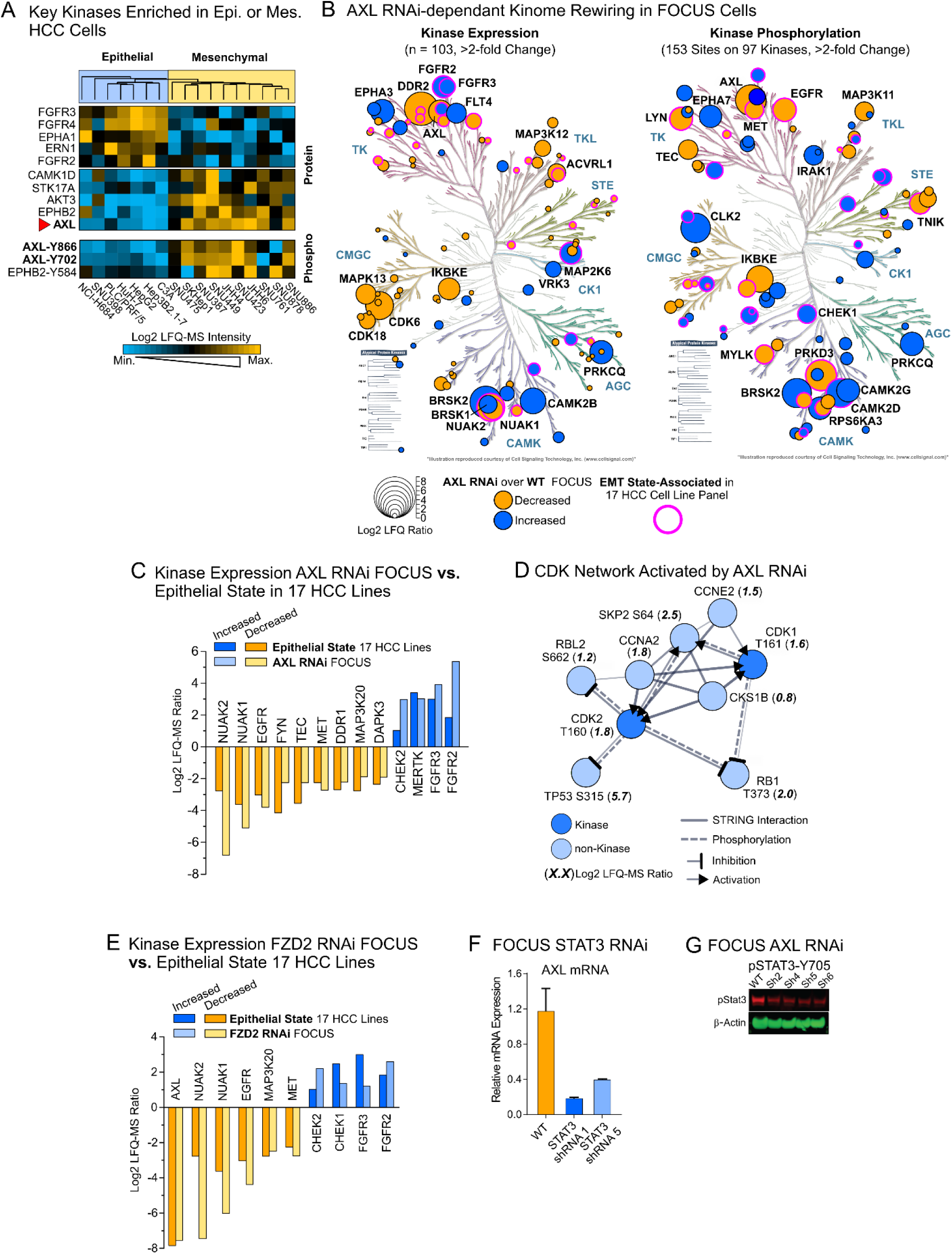
AXL regulates kinase signaling in HCC cell EMT and is a transcriptional target of FZD2-FYN/STAT3. (A) Relative expression of the top 5 kinases associated with either the mesenchymal or epithelial phenotype across 17 HCC cell lines. Quantified activating sites on AXL and EPHB2 are shown. (B) Kinase protein (left) and phosphosite (right) expression differences in FOCUS *AXL* RNAi cells over FOCUS WT cells overlaid on the human kinome dendrogram. The top 20 highest regulated kinases are labeled. The phosphorylation site with the highest absolute LFQ ratio on each kinase was used for plotting. (C) Highly regulated kinases from *AXL* knockdown in FOCUS cells also found associated with EMT in the larger panel. (D) STRING interaction network of CDK1 and 2 substrates and corresponding CDK1/2 phosphorylation sites enriched in *AXL* RNAi FOCUS cells over WT. (E) Kinases regulated by *FZD2* RNAi in FOCUS cells that were also associated with EMT. (F) qPCR of *AXL* mRNA in FOCUS STAT3 RNAi cells. (G) Western blot of pSTAT3-Y705 in FOCUS *AXL* RNAi cells. See also Figure S5 and **Table S6**.

Consequently, we used RNAi to knock down *AXL* in the FOCUS cell line, a widely used mesenchymal HCC model to study the EMT (Figure S5A) (*40*). Western blot analysis of four EMT marker proteins and qPCR quantification of 43 EMT marker mRNAs confirmed that *AXL* RNAi induced widespread expression changes indicating mesenchymal-epithelial transition (MET, Figure S5B-D). Next, we compared the FOCUS *AXL* RNAi line to the wild-type control using LFQ kinobead/LC-MS and found >2-fold expression changes in 103 protein kinases and 153 phosphosites on 97 kinases (Figure 5B, **Table S6**). Importantly, AXL-regulated kinase expression in FOCUS cells overlapped greatly with the EMT state-associated kinome in the 17 HCC cell lines (Figures 5B, S5E and **S5F**). We found, for example, that expression of the mesenchymal kinases NUAK1 and 2, FYN, EGFR, and the cabozantinib target c-MET decreased >4-fold in response to *AXL* knockdown. Concurrently, kinases associated with the epithelial state such as FGFR2/3 and CHEK2 increased in expression in *AXL* RNAi cells (Figure 5C and **Table S6**). Notably, epithelial FOCUS *AXL* RNAi cells showed the same increased activity of cell cycle-related signaling components that we found in the epithelial cells of our 17 HCC line panel. These signaling components included a network of nine nodes centered around phospho-activated CDK1 and 2, including key phosphorylation sites and interactions for CDK1/2 and their substrates TP53, RB1, RBL2, and CCNA2 (Figure 5D). Collectively, these results highlight the effects of AXL in HCC cell EMT and identify downstream kinases that may play important roles in the EMT and drug response phenotype.

### FZD2 is a master regulator of AXL expression and EMT-associated kinome rewiring

Having identified various kinases downstream of AXL, we next sought to identify upstream signaling components that could regulate AXL expression and the EMT in HCC cells. We found previously that *AXL* mRNA co-expresses with *FZD2* mRNA and showed here that this relationship is conserved in patient HCCs (Figure 4D). FZD2 is a G-protein coupled receptor for Wnt5A/B and regulates HCC cell EMT via a FYN/STAT3-dependent pathway (*40*). To investigate a possible functional connection between FZD2, AXL, and other downstream pathway components, we profiled our FOCUS *FZD2* RNAi cell model with LFQ kinobead/LC-MS (Figure S5G and S5H, **Table S6**). Indeed, expression of AXL and 86 other kinases downstream of AXL was significantly affected by *FZD2* knockdown (ratio >2-fold, Figure 5E and S5G), suggesting the existence of a FZD2-AXL pathway important for EMT state-associated kinome rewiring in HCC. and other downstream pathway components, we profiled our FOCUS *FZD2* RNAi cell model with LFQ kinobead/LC-MS (Figure S5G and S5H, **Table S6**). Indeed, expression of AXL and 86 other kinases downstream of AXL was significantly affected by *FZD2* knockdown (ratio >2-fold, Figure 5E and S5G), suggesting the existence of a FZD2-AXL pathway important for EMT state-associated kinome rewiring in HCC. To investigate if AXL expression is connected to FZD2’s activation of FYN and STAT3, we knocked down *STAT3* in FOCUS cells and found that *AXL* mRNA levels dropped drastically (Figure 5F); these results implicate that in HCC cells AXL may be a transcriptional target of STAT3, further supporting the existence of a FZD2-FYN/STAT3-AXL signaling module. We also found that *AXL* RNAi in FOCUS cells affects the phosphorylation of STAT3 at its activating site, pY705, identifying a probable feed-forward loop from AXL to STAT3 that reinforces AXL expression (Figure 5G). These results integrate AXL into the greater framework of FZD2-regulated HCC cell EMT and reveal additional kinase targets for pharmacological intervention.

### NUAK1 and 2 stabilize AXL expression and promote the EMT in HCC

To identify novel drug targets to reverse EMT and overcome drug resistance, we next looked for kinases downstream of FZD2-FYN/STAT3-AXL signaling module. The nuclear serine/threonine kinases NUAK1 and NUAK2 were tightly associated with the mesenchymal state in the 17 HCC lines and their expression greatly decreased with *AXL* and *FZD2* RNAi (20 to 30-fold, Figure 5C and 5E). Furthermore, NUAK1 is implicated in tumor metastasis (*41*) and both NUAK1 and NUAK2 were shown to promote tumor cell survival (*42, 43*). We reasoned that NUAK1 and NUAK2 are top candidates for promoting the HCC cell EMT and drug resistance.

To test if NUAK1 and NUAK2 are *bona fide* drivers of the HCC cell EMT or only bystanders, we generated stable FOCUS *NUAK1* or *NUAK2* RNAi cell lines (Figure 6A). Remarkably, knockdown reduced cell migration by 60-70%, compared to a 25% reduction in FOCUS *AXL* RNAi cells (Figure 6B and C), suggesting a prominent role of NUAK1 and 2 in HCC cell EMT. Next, we compared FOCUS *NUAK1* and *NUAK2* RNAi cells to WT cells using LFQ kinobead/LC-MS and found that *NUAK1* or *NUAK2* knockdown affected the expression of 92 and 97 protein kinases >2-fold, respectively (Figure S6A, **Table S6**). Surprisingly, the kinome profiles of *NUAK1* and *NUAK2* RNAi cells were very similar. The expression of 43 out of 52 highly regulated kinases (LFQ-MS ratio >4-fold) was affected in both knockdown lines and the LFQ-MS ratios of all significantly regulated kinases had a Pearson’s r value of 0.91 (Figure 6D and **Table S6**).

**Figure 6.**
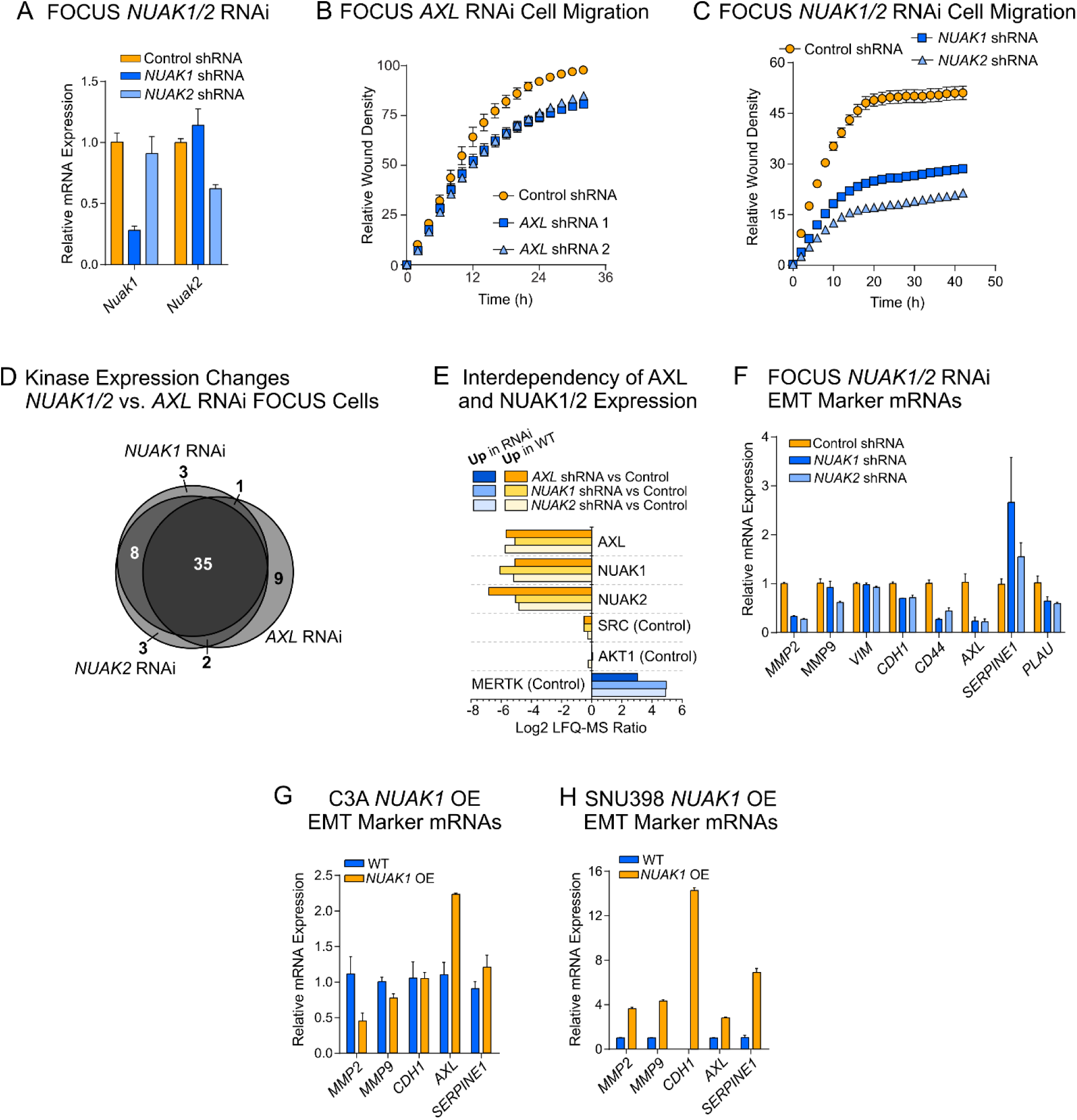
Major role of NUAK1/2 in HCC cell EMT through a FZD2-AXL-NUAK signaling module. (A) qPCR of *NUAK1* and *NUAK2* mRNA in FOCUS *NUAK* RNAi cells. (B) Wound healing assay in FOCUS *AXL* RNAi cells compared to control shRNA cells. (C) Wound healing assay in FOCUS *NUAK1* and *NUAK2* RNAi cells compared to controls. (D) Venn diagram of effect of *NUAK1* RNAi, *NUAK2* RNAi and *AXL* RNAi on kinase expression in FOCUS cells. Only kinases with an LFQ-MS expression change >4-fold are shown. (E) Change of protein expression of AXL, NUAK1, and NUAK2 with three kinases (AKT1, SRC and MERTK) as internal controls upon knockdown of either of these kinases. (F) qPCR of mRNA changes in EMT markers upon knockdown of either *NUAK1* or *NUAK2* in FOCUS cells. (G, H) Effect of *NUAK1* overexpression (OE) on EMT markers in C3A and SNU398 cells, respectively. See also Figure S6 and **Table S6**.

Strikingly, when we compared *AXL* RNAi-regulated kinase expression with *NUAK1* and *NUAK2* RNAi-regulated kinase expression, 85% of highly regulated kinases (LFQ-MS ratio >4-fold) were common between *AXL, NUAK1* or *NUAK2* RNAi experiments and the LFQ-MS ratios of all regulated kinases showed a r-value of 0.93 (Figure 6D). Furthermore, AXL expression levels were greatly decreased by either *NUAK1* or *NUAK2* RNAi and *vice versa*. This hinted at the existence of a positive feedback mechanism where NUAK kinases regulate AXL expression or protein stability. Indeed, our qPCR analysis of EMT markers confirmed that *AXL*, *CD44* and *MMP2* were decreased in FOCUS *NUAK* RNAi cells (Figure 6F). Additionally, ectopic expression of *NUAK1* in epithelial-type C3A and SNU398 cells caused at least partial EMT as indicated by increased expression of *MMP2* and *MMP9*, *AXL* and *SERPINE1* (Figure 6G, H and S6B). These results confirm the involvement of NUAK1 and NUAK2 in HCC cell EMT and their tight functional connection with AXL suggests that they could be surrogate targets for AXL in the development of new EMT-targeted therapies. We hypothesized, therefore, that modulating NUAK1 and NUAK2 could potentially reverse EMT and restore sensitivity to kinase inhibitors in mesenchymal HCC cells.

### AXL-NUAK1/2 signaling protects HCC cells against replication stress and promotes resistance to cell cycle checkpoint inhibitors

We discovered a tight functional connection between the important EMT kinase AXL and NUAK1 and NUAK2. These kinases are significantly overexpressed in drug resistant mesenchymal HCC lines and their knockdown by RNAi reverses the EMT. We thus reasoned that the combination of either AXL or NUAK inhibitors with drugs that kill epithelial HCC cells could be an effective approach to kill drug-resistant mesenchymal cells. To identify specific vulnerabilities and suitable candidate drugs, we examined pathways that became activated by the MET in greater detail.

When we knocked down FZD2-AXL-NUAK pathway components in FOCUS cells, we observed increased expression of the FGFR and activation of mitogenic signaling and cell cycle cues. We also found increased cell cycle activity in the epithelial fraction of the 17 HCC line panel. Interestingly, this increased cell cycle activity was accompanied by elevated activity of kinases known to regulate the DNA damage response and cell survival under replication stress conditions (44). For instance, we observed phospho-activation of CHEK1 and 2 along with increased phosphorylation of CDK2, ERK1/2, ATR, CHEK1/2 and WEE1 substrate sites (Figure 7A, S6C and S6D). Similarly, AXL RNAi in the FOCUS line activated the DDR kinases CHEK1, WEE1 and increased phosphorylation of their substrates CDK11B and CDK2 (Figure 7B and S6E). These findings suggest that AXL, NUAK1 and NUAK2 suppress the cell cycle and DNA damage response signaling, and protect HCC cells from replication stress. Importantly, we also observed that the efficacy of KI drugs that target cell cycle-related kinases was strongly reduced in the mesenchymal fraction of our 17 HCC line panel (Figure 7C). Among the inhibitor classes, the efficacy of CDK, PLK1 and CHEK1/2 inhibitors (KI cluster 9) were the most dependent on the target cell’s EMT state (Figure 7C and S2A). This suggests that the survival of epithelial HCC cells may depend on DNA damage checkpoint signaling (*44*), and we hypothesized that combinatorial inhibition of AXL, NUAK1 and NUAK2, and cell cycle checkpoint kinases could efficiently kill mesenchymal HCC cells (Figure 7D) (*45*).

**Figure 7.**
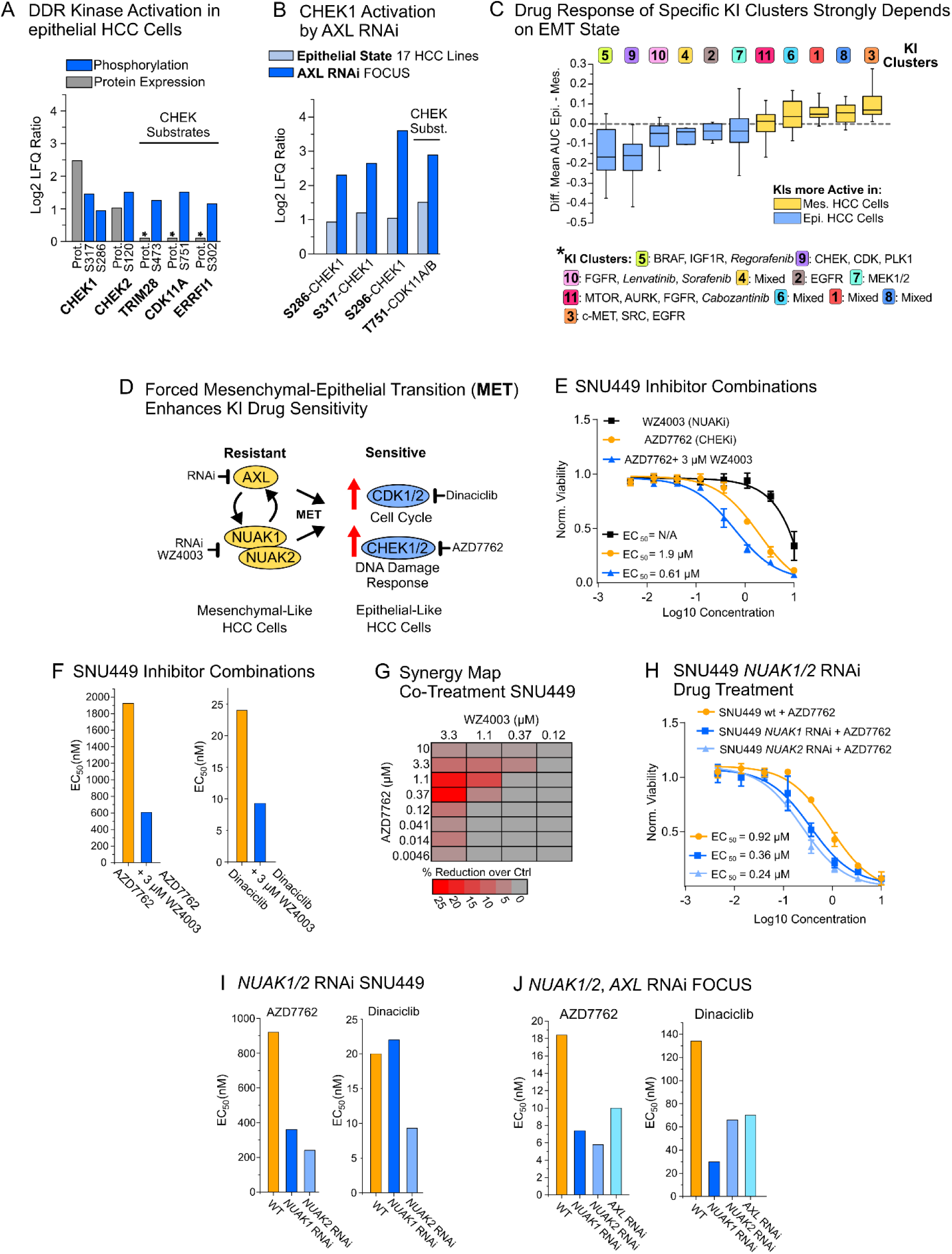
Perturbation of AXL or NUAK1/2 function induced MET and increases HCC cell sensitivity to cell cycle- and DNA damage checkpoint-kinase inhibitors. (A) Activating phosphosites on CHEK1 and 2 and their substrates enriched in 7 epithelial vs. 10 mesenchymal HCC cell lines. (B) Phosphorylation sites on CHEK1 and its substrates that indicate activation of this kinase in FOCUS *AXL* RNAi cells over WT and that are associated with EMT in the 17-cell line panel. (C) Difference of mean AUC values in epithelial vs. mesenchymal HCC cells for the 299 KI panel subdivided by membership in the 11 KI pathway clusters (see Figure 2A and **Table S3**). (D) Strategy inducing MET and drug sensitivity through NUAK1/2 and AXL inhibition. (E) and (F) EC_50_-curves and bar plots of drug co-treatment experiments in SNU449 cells (n = 4, error bars are S.D.) (G) Heatmap of drug synergy and titratability of AZD7762 and WZ4003 in SNU449 cells. (H) and (I) EC_50_-curves and bar plots of drug treatment experiments with AZD7762 and dinaciclib in the SNU449 *NUAK1/2*-RNAi cell lines. (J) Drug response to AZD7762 and dinaciclib in FOCUS *NUAK1/2* and *AXL* RNAi cell lines. See also Figure S6

To test this strategy, we cotreated the mesenchymal and drug resistant SNU449 line (**Table S1** and **S3**) with the selective NUAK1 and NUAK2 inhibitor WZ4003 (*46*) to induce MET and the checkpoint kinase inhibitor AZD7762 to kill these cells. Additionally, we used the CDK inhibitor dinaciclib to test if elevated MET-dependent cell cycle activity translated into increased efficacy of such drugs. Remarkably, cotreatment resulted in a reproducible decrease in EC_50_s, i.e. ∼three-fold and two-fold for both CHEK and CDK inhibitors (Figure 7E and 7F). This effect was dose-dependent for both AZD7762 and WZ4003 in SNU449 cells, and still significant at 1 µM of WZ4003 (Figure 7G). These encouraging results led us to establish SNU449 *NUAK1* or *NUAK2* RNAi cells (Figure S6F). Gratifyingly, testing of SNU449 *NUAK* RNAi cells and WT controls with varying doses of AZD7762 and dinaciclib recapitulated the drug co-treatment results (3-4-fold decrease in EC_50_s, respectively, Figure 7H and 7I). To consolidate our findings, we next tested our FOCUS *NUAK* and *AXL* RNAi cell lines against AZD7762 and dinaciclib. As in SNU449 cells, *NUAK1/2* RNAi FOCUS cells were sensitized to kinase inhibition up to 4-fold (Figure 7J). Together, these results establish, for the first time, that NUAK1 and NUAK2 can regulate HCC cell resistance to targeted therapy, which has important implications the design of novel cancer therapeutics.

## DISCUSSION

We have described a novel pharmacoproteomics platform that combines comprehensive kinome profiling (*10*) and KI drug screening to identify signaling pathways associated with cancer drug sensitivity and resistance. Our kinobead/LC-MS profiling technology determines not only kinase expression and their phosphorylation states, but also kinase-dependent signaling complexes and their phosphorylation (*47*), therefore greatly improving analytical coverage of the kinome and kinase-dependent signaling networks over conventional global phosphoproteomics methods (*10*).

Hepatocellular carcinoma (HCC) is a deadly cancer with no known druggable genetic alterations; because patient response rates to FDA-approved HCC drug remain low (10-15%) there is a pressing need to discover novel predictive biomarkers and drug targets (*15*). We profiled 17 HCC cell lines with our kinobead/LC-MS technology and correlated kinome activity with drug response, creating a map of molecular markers for HCC cell sensitivity and resistance to 299 KIs, including the four clinical HCC drugs sorafenib, regorafenib, lenvatinib and cabozantinib (*16–18, 48*). Remarkably, our results show that the phosphorylation and thus the activation states of kinases are better predictors of drug response than mRNA and protein expression levels. Hence, precision oncology must integrate genomic and transcriptomic data with information on protein expression and, most importantly, protein PTMs to discover optimal biomarkers and drug targets (*5*). Even more importantly, we leveraged our dataset of kinome features to identify pathway-based markers of drug response. Such biomarkers typically outperform ones that are based on single molecular features (*28, 29*) and our dataset includes a comprehensive collection of pathway-based biomarkers that should serve as an important resource for oncology research. In support of this notion, we already demonstrated that our kinobead/LC-MS platform can determine kinome activity in individual patient HCCs, and that our pathway-based biomarkers are enriched in these tumors.

Notably, clustering of the 299 KIs by the similarity of the GSEA-enriched pathways also allowed us to classify KIs by drug action instead of literature-based annotation of their targets. We speculate, therefore, that akin to pharmacogenomics efforts in cancer cell lines (*3, 4*), our pharmacoproteomics platform can identify *bona fide* KI targets as well as uncover mechanisms of action for KIs with unspecified polypharmacology.

Finally, our analyses of HCC drug response pathways identified the EMT as an important mechanism of drug resistance for a broad range of KI drugs (*24*). With the EMT as a classifier, we identified 101 kinases that are highly active in either epithelial or mesenchymal HCC cells, including well characterized EMT kinases such as AXL, c-Met and FYN, and poorly characterized kinases like NUAK1 and 2, CAMK1D, EPHB2, and many others. Hence, our dataset represents the first comprehensive analysis of kinase-dependent signaling networks that control cancer cell EMT. Mining of our dataset and additional functional studies revealed a FZD2-AXL-NUAK1/2 signaling module that is highly active in mesenchymal HCC cells. Remarkably, when we inhibited either NUAK1, NUAK2 or AXL by RNAi knockdown or pharmacological inhibition mesenchymal HCC cells underwent MET, in turn activating targets of clinical and preclinical HCC drugs such as the FGFR and the cell cycle (*17, 33*). Confoundingly, this MET-dependent increase in cell cycle activity was accompanied by induction of DNA damage response signaling, indicating that rapidly dividing epithelial HCC cells suffer from near-catastrophic replication stress (*44*). Based on these insights, we developed a strategy to treat mesenchymal-type drug resistant HCC: we first induce MET by inhibiting kinases that stabilize the mesenchymal phenotype, and then kill these cells by treatment with a cell cycle checkpoint kinase inhibitor. We showed here that inhibition of either NUAK1 and 2, or AXL leads to enhanced cell killing with CHEK and CDK inhibitors, and we speculate that our strategy can be expanded to other EMT kinases and HCC drugs.

In summary, our pharmacoproteomics platform discovers vulnerabilities in relevant cancer models and identifies rational drug combinations and new therapeutics. Reproducible and amenable to lab automation, we expect that our pharmacoproteomics platform can be easily scaled up to profile hundreds of clinical tumor samples or cell lines. Our platform will be useful to screen kinome activity and drugs in tumor-derived organotypic slice cultures (*49*) or organoid cultures (*50*), discovering markers of drug response, classify drug-sensitive disease subtypes, and novel drug targets in these advanced cancer models. Importantly, our study provides the first comprehensive map of the kinase-dependent signaling networks that define the mesenchymal and epithelial state in cancer; our quantitative measurements of kinome abundance and phosphorylation sites significantly expands our knowledge of EMT mechanisms. Our dataset of kinome features and signaling pathways and their relevance to drug effects for 299 KIs likely applies to other cancer cell types and is an important resource for researchers studying kinase inhibitors and kinase-dependent cell signaling.

## Supporting information

Log2 mRNA intensity of 50 EMT and stem cell markers as well as 509 protein kinases in the 28 CCLE HCC cell lines. Names and primary targets of the 299

MaxQuant protein groups and phospho(S/T/Y) output data obtained from kinome profiling of 17 HCC cell lines. Missing values were replaced by data imput

Results of the HTS of 299 kinase inhibitors in 17 HCC cell lines. Summary table of AUC values. Summary of Reactome pathways used in the kinome-GSEA an

Results from Pearson correlation between mean MS intensity values of all quantified proteomics features (n = 13935) and kinase inhibitor AUCs (n = 299

Results of protein kinase quantification by super SILAC kinobead/LC-MS profiling in 9 paired (tumor/NAL) human HCC specimens. MaxQuant protein groups

MaxQuant protein groups and phospho(S/T/Y) output data based on kinome profiling of the FOCUS cell AXL, FZD2 and NUAK1/2 RNAi models.

## ACKNOWLEDGMENTS

We thank Dr. Timothy J. Martins and James Annis from the Quellos HTS Core at ISCRM of the University of Washington for providing kinase inhibitor HTS services. We thank John D Scott, Anthony K. Leung, David Shechner, and members of the Ong and Maly Labs for comments on the manuscript. This work was supported by grants from the National Institutes of Health issued under the award numbers R01GM086858, R01GM129090, R01AR065459, R21EB018384, R21CA177402, K22CA201229-01, by the Sidney Kimmel Foundation and by a postdoctoral research fellowship of the German Research Foundation (DFG) awarded to M.G. (GO 2358/1-1). We gratefully acknowledge funding from the Department of Defense under the award number CA150370 as well as the Fibrolamellar Cancer Foundation. This work used an EASY-nLC1200 UHPLC and Thermo Scientific Orbitrap Fusion Lumos Tribrid mass spectrometer purchased with funding from a National Institutes of Health SIG grant S10OD021502 (S-E.O.) The content is solely the responsibility of the authors and does not necessarily represent the official views of the National Institutes of Health.

## AUTHOR CONTRIBUTIONS

Conceptualization, M.G., S-E.O. and T.S.G; Methodology, M.G; Investigation, M.G., M.C., A.S. and V.N.V.; Formal Analysis, M.G., H-T.L.; Writing – Original Draft, M.G., S-E.O.; Writing – Review and Editing, M.G., D.J.M, T.S.G, R.S.Y, S-E.O.; Funding Acquisition, S-E.O., D.J.M, T.S.G and R.S.Y; Supervision, S-E.O.

## DECLARATION OF INTERESTS

The authors declare that there are no competing financial interests.

## SUPPLEMENTARY FIGURES, TITLES, AND LEGENDS

**Figure S1.**
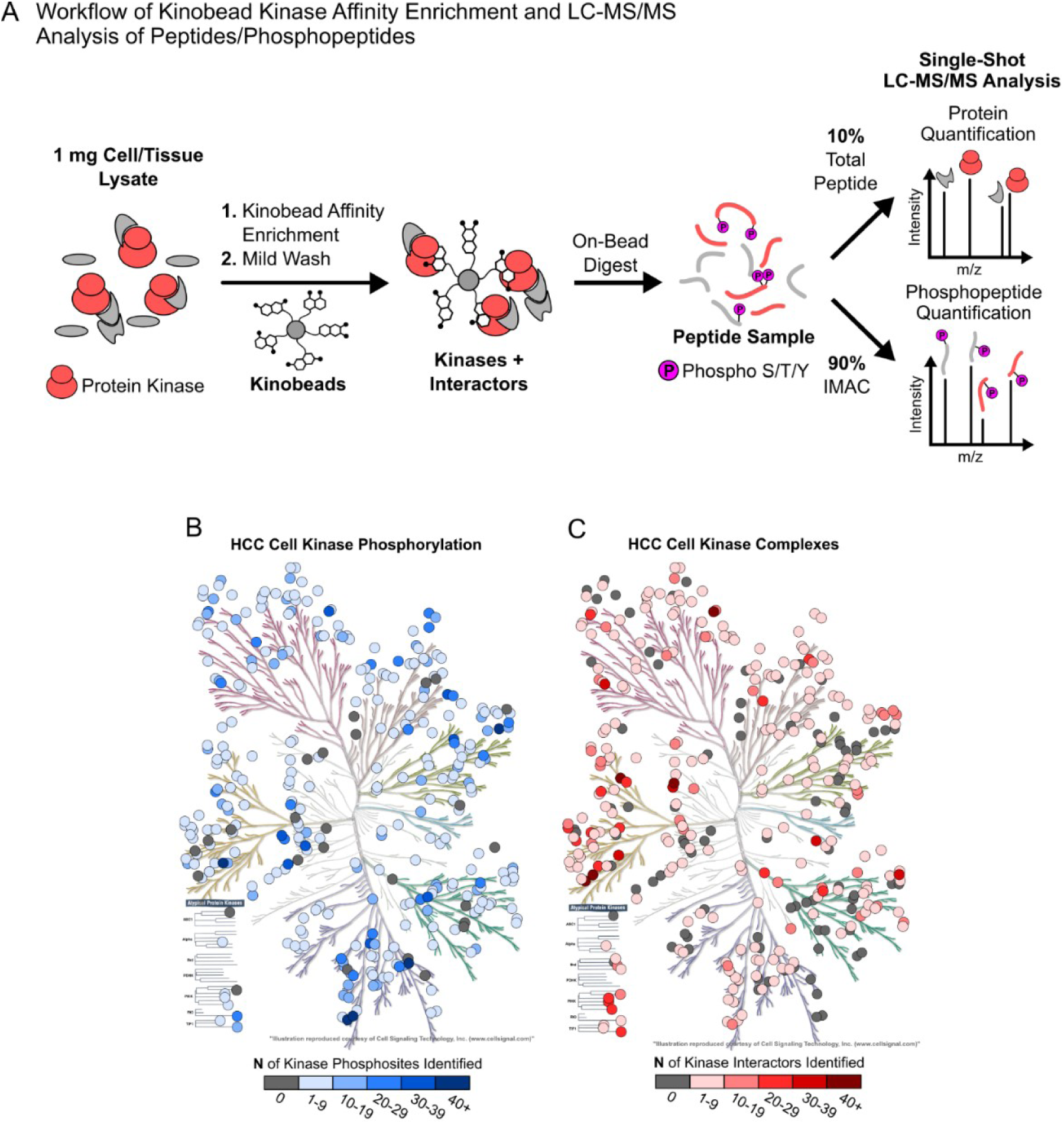
Kinobead/LC-MS workflow and kinase, their phosphorylation sites and complex components identified in the 17 HCC cell line panel, Related to Figure 1. (A) Kinobead/LC-MS workflow. (B) The human kinome dendrogram overlaid with quantified kinases (n = 346, circles) and number of phosphosites (color scale) for each kinase. (C) As in (B) but color scale indicates the number of quantified interactors for each kinase. This includes only BioGRID interactions with at least two lines of experimental evidence (see ‘Materials and Methods’).

**Figure S2.**
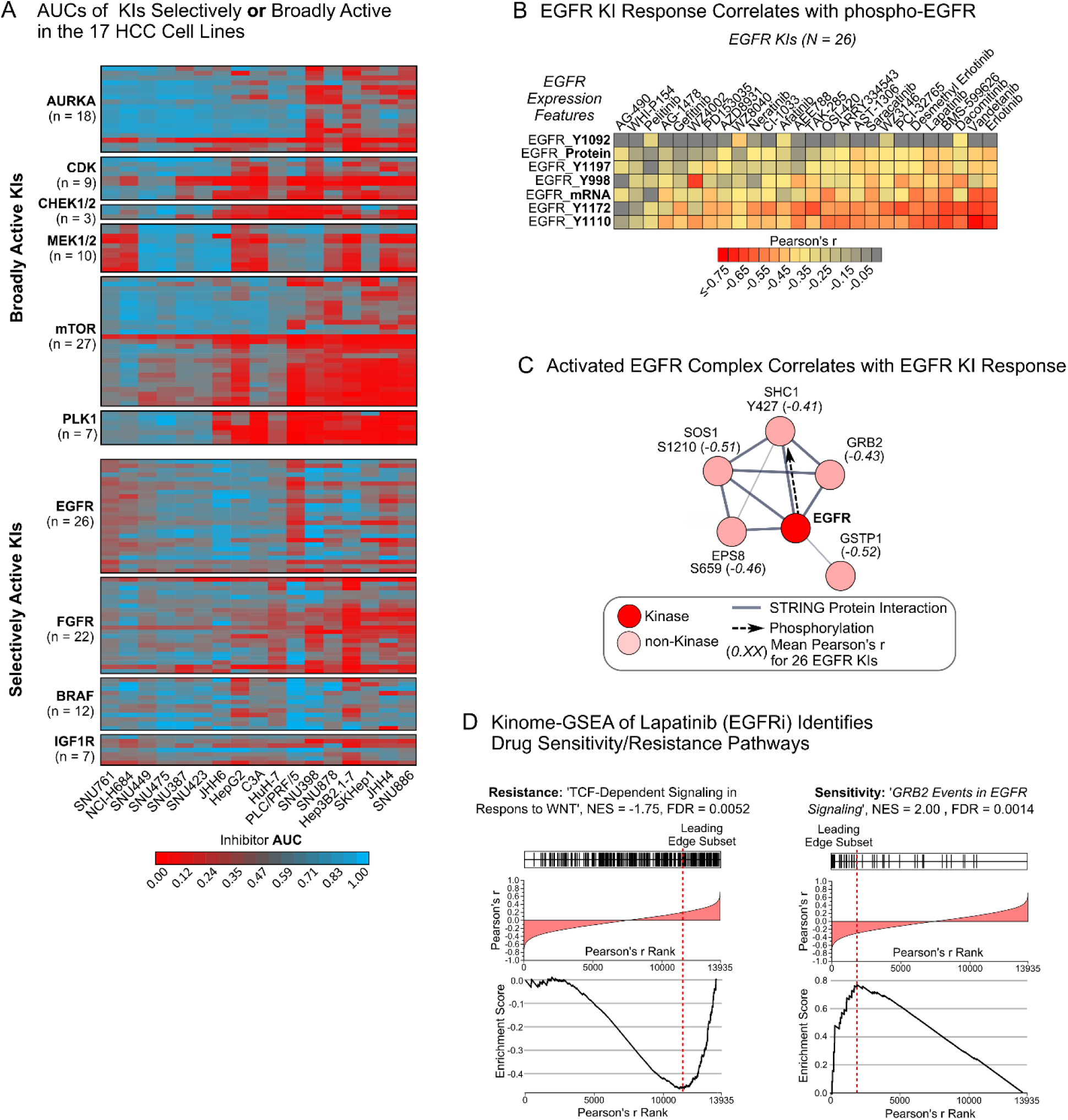
HTS drug screen results, AUC-kinome feature correlation and kinome-GSEA. Related to Figure 1. (A) KI AUC values across the 17 HCC cell line panel. Low AUCs correspond to a strong drug response, i.e. more cell killing. Representative broadly active KIs (AURK, CDK, CHEK1/2, MEK1/2, MTOR, PLK1) are shown in the top panel. Representative KIs active only in specific cell lines (EGFR, FGFR, BRAF and IGF1R) are shown in the bottom panel. (B) Heatmap of Pearson’s r-values from correlation of EGFR expression features (mRNA, protein, and phosphosites) with the AUC values of 26 EGFR inhibitors (see ‘Materials and Methods’ and **Table S3**). (C) STRING network (*51*) of proteins interacting with the activated EGFR. Mean r-values for proteomics feature–AUC correlation are shown for 26 EGFR inhibitors (**Table S3**).

**Figure S3.**
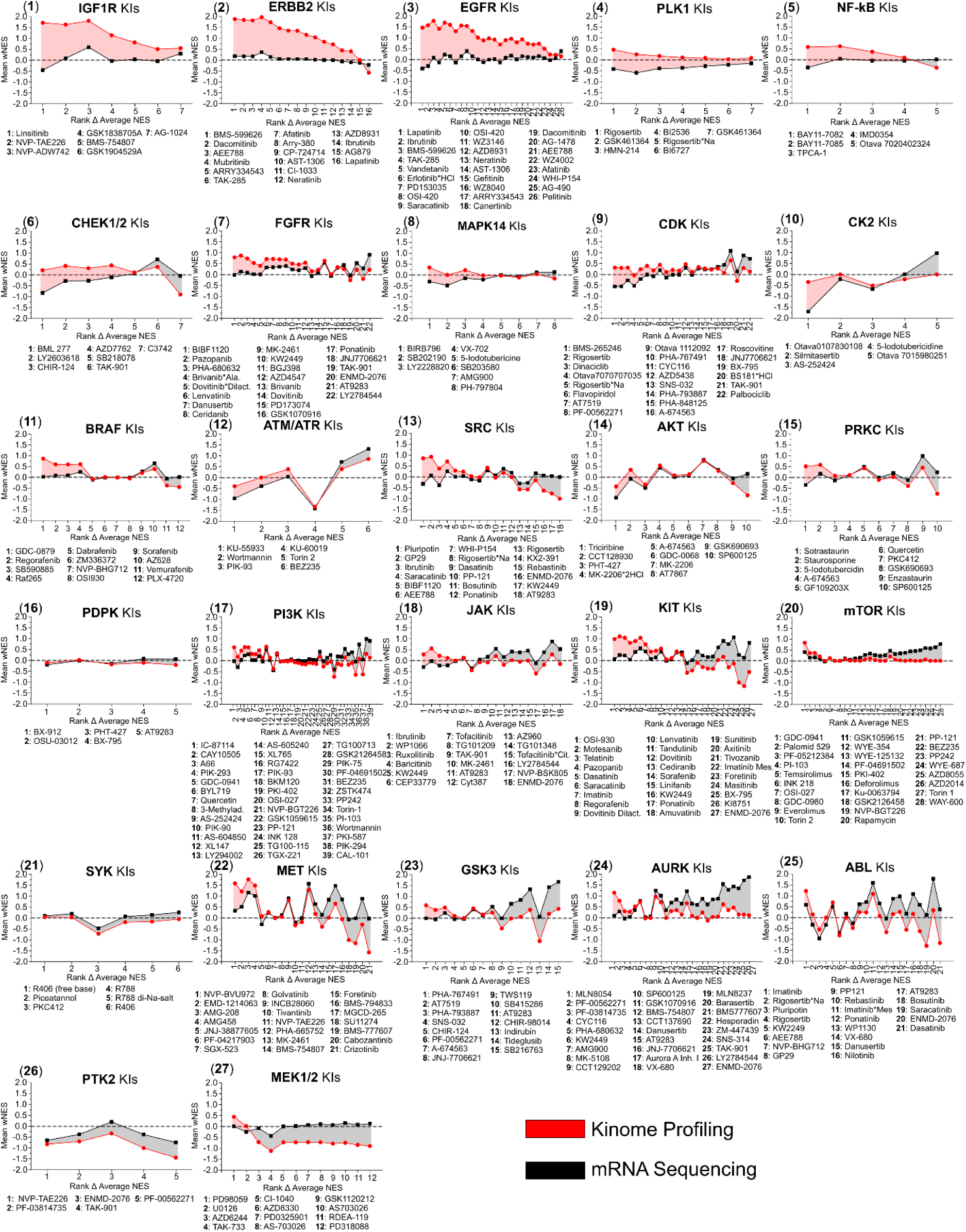
Comparison of kinome-GSEA and mRNA-GSEA in predicting target pathways of KIs. For each KI in the screening panel (*N* = 299) mean wNES values (wNES = NES*(1-FDR)) were calculated across all Reactome pathways of which the main target of the KI is a member. This was done for both, kinome-GSEA and mRNA analysis (see ‘Materials and Methods’) and compounds were grouped by their main target (see **Table S3**). wNES were in each graph are sorted by difference wNES (kinome) – wNES (mRNA).

**Figure S4.**
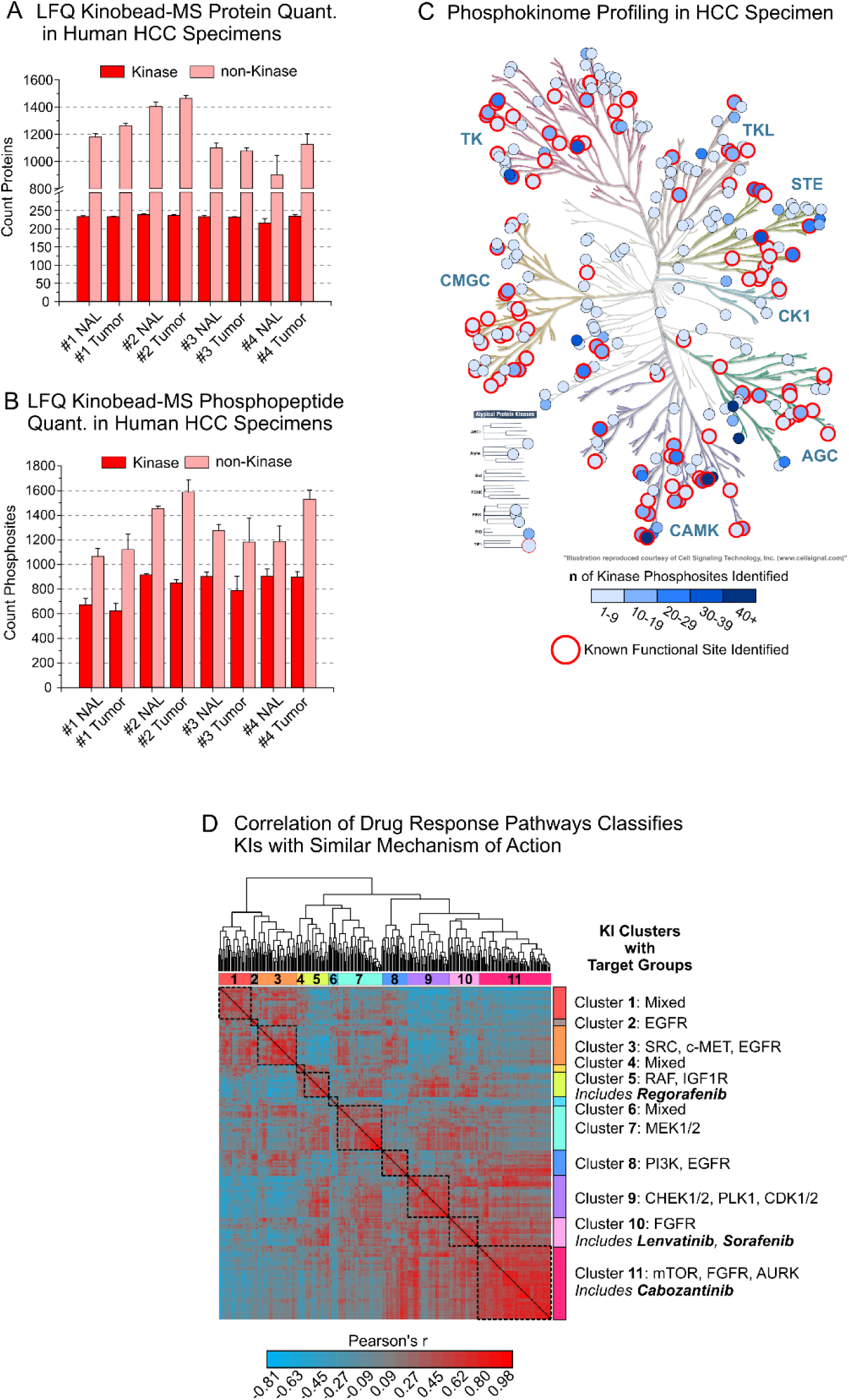
Kinobead/LC-MS profiling of clinical HCC specimens and KI clustering by kinome-GSEA enrichment scores. (A) Mean number of kinases and other protein quantified in single LFQ kinobead/LC-MS runs from human HCC and NAL tissue specimens. Error bars are the S.D., n = 6. (B) Same as in (A) but phosphopeptides derived from kinases and other proteins. (C) Kinases and identified phosphopeptides in human HCC specimens. Kinases for which known functional phosphopeptides were identified are highlighted in red (see also (A)). (D) Pairwise correlation of GSEA Reactome pathway scores (FDR-weighted NES, wNES) for 299 KIs; r-values are clustered into 11 groups of KIs by their similarity in enriched signaling pathways. Partial lists of primary targets (from literature) within KI clusters are shown. The complete list is available in the interactive web application (*25*).

**Figure S5.**
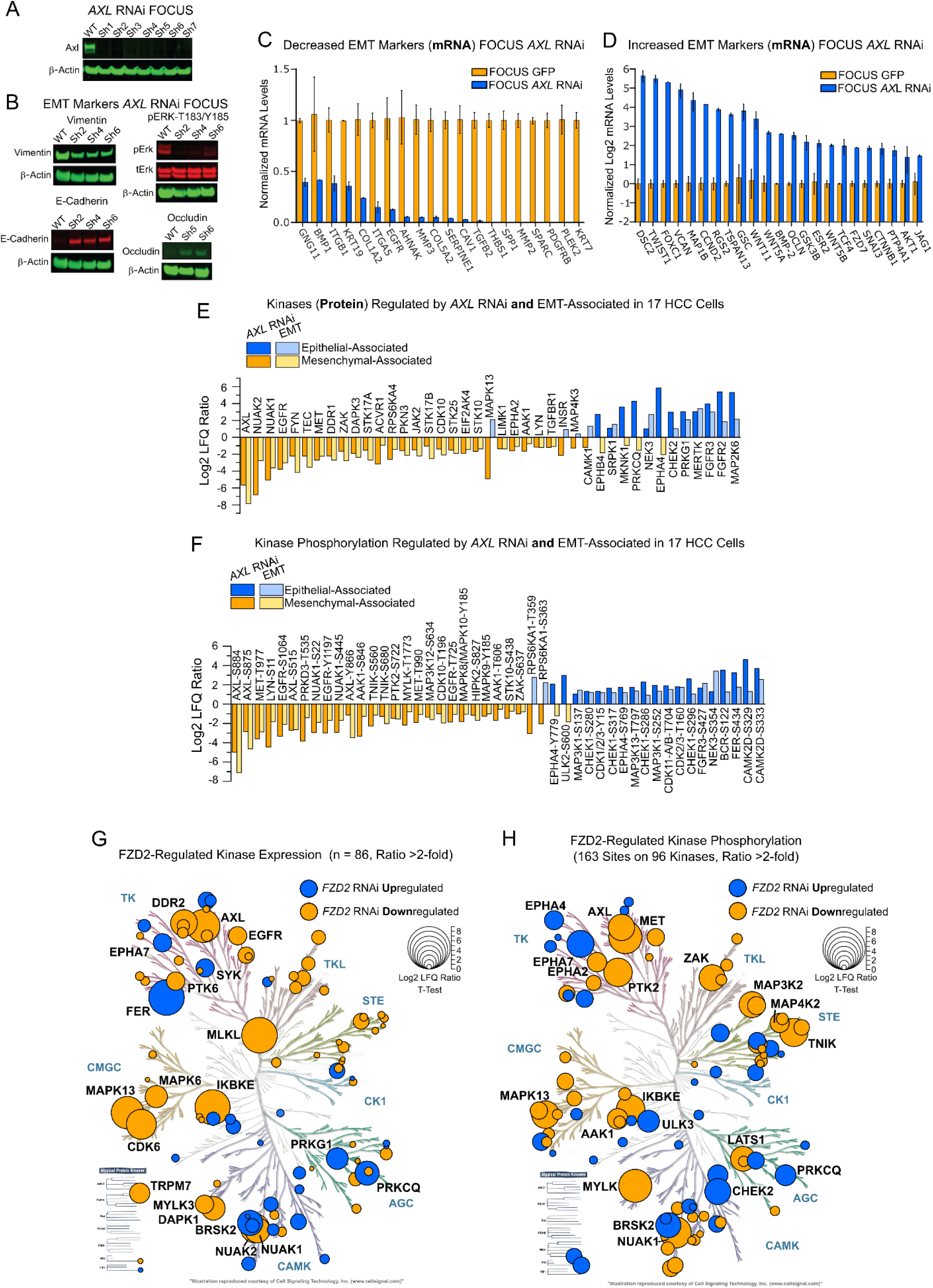
AXL regulates kinase signaling in HCC cell EMT and is a transcriptional target of FZD2-FYN/STAT3, Related to Figure 5. (A) Western blot analysis of AXL in FOCUS *AXL* RNAi cells. (B) Western blot analysis of FOCUS *AXL* RNAi cells and FOCUS WT cells for expression of the EMT markers vimentin, E-cadherin, occludin and phospho ERK. (C) EMT marker mRNAs that were increased in expression upon *AXL* RNAi knockdown in FOCUS cells as quantified by qPCR. (D) EMT marker mRNAs that were decreased in expression upon *AXL* RNAi knockdown in FOCUS cells as quantified by qPCR. (E) Kinases that changed in expression >2-fold in FOCUS *AXL* RNAi cells over FOCUS WT cells that were also significantly associated with EMT in the 17 HCC cell line panel. Kinase expression was quantified using kinobead/LC-MS in six biological replicate experiments with LFQ. Bars are the difference of the mean MS Intensity (LFQ ratio) from a two-tailed Student’s T-test, Benjamin-Hochberg FDR = 0.05). (F) Kinase phosphopeptides that changed in expression >2-fold in FOCUS *AXL* RNAi cells over FOCUS WT cells that were also significantly associated with EMT in the 17 HCC cell line panel (see (E)). (G) FOCUS *FZD2* RNAi-regulated kinase expression quantified by kinobead/LC-MS. Filled circles represent individual kinases and the circle size scales with log2 LFQ ratio relative to control. Only kinases with a >2-fold expression change are shown. Statistically significant kinase regulation was determined with a two-tailed Student’s T-test with BH correction for multiple comparison (FDR = 0.05, n = 6 in each state). (H) FOCUS *FZD2* RNAi-regulated phosphorylation expression (see (G)). The highest regulated phosphorylation site on each kinase (log2 ratio) was used for plotting circle size.

**Figure S6.**
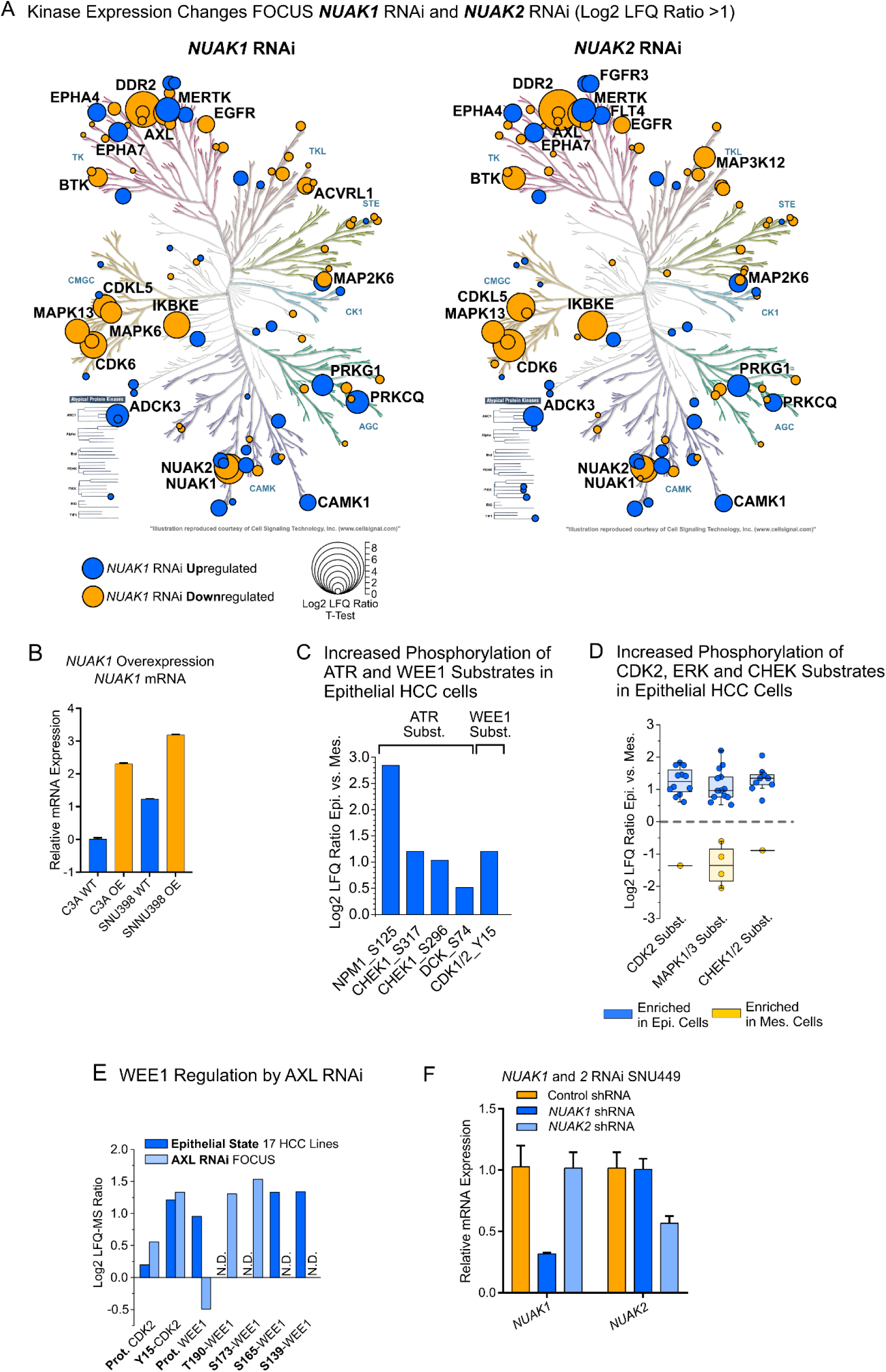
Major role of NUAK1/2 in HCC cell EMT through a FZD2-AXL-NUAK signaling pathway, Related to Figure 6 and 7. (A) FOCUS *NUAK1* RNAi-regulated kinase expression quantified by kinobead/LC-MS. Filled circles represent individual kinases and circle size scales with the log2 LFQ ratio relative to control. Only kinases with a >2-fold expression change are shown. The kinase’s phosphosite ratio with the highest magnitude between drug response phenotypes is plotted. Statistically significant kinase regulation was determined with a two-tailed Student’s T-test with BH correction for multiple comparison (FDR = 0.05, n = 6 in each state). (B) qPCR analysis of *NUAK1* mRNA in C3A and SNU398 cells overexpressing *NUAK1* plasmid DNA. (C) Difference in expression of known ATR and WEE1 phospho-substrate sites (Phosphosite Plus) between epithelial and mesenchymal HCC cells. (D) Difference in expression of known CDK2, MAPK1/3 and CHEK1/2 phospho-substrate sites (Phosphosite Plus) between epithelial and mesenchymal HCC cells. (E) Phosphorylation sites on WEE1 and its substrates that indicate activation of this kinase in FOCUS *AXL* RNAi cells over WT and that are associated with EMT in the 17-cell line panel. (F) qPCR analysis of *NUAK1* and *NUAK2* mRNA in *NUAK1* and *2* RNAi SNU449 cells, respectively.

## MATERIALS AND METHODS

### Contact for reagents and resources sharing

As lead contact, Shao-En Ong is responsible for all reagent and resource requests. Please contact Shao-En Ong at with requests and inquiries.

#### Experimental model and subject details

##### Cell lines and cell culture conditions

C3A, HepG2, SNU398, PLC/PRF/5, Hep3B2.1-7, SKHep1, SNU475, SNU387, SNU423, SNU449, HCT-116, SH-SY5Y, U-2 OS, Du4475, HL-60, Jurkat and K562 cell lines were purchased from the American Type Culture Collection (ATCC). NCI-H684, SNU761, SNU886 and SNU878 were purchased from the Korean Cell Line Bank (KCLB). JHH4, JHH7 and HuH-7 cells were purchased form the JRCB Cell Bank. FOCUS cells were obtained from the Laboratory of J. Wands, Brown University (*52*). All cells were grown at 37°C under 5% CO2, 95% ambient atmosphere. A large number of cryofrozen cell stocks were generated from the original vial from the cell bank. Experiments were performed with cells at <10 passages from the original vial. All cell media used were supplemented with 100x penicillin-streptomycin-glutamine (Thermo Fisher Scientific, Waltham, MA). FOCUS and HuH-7 cells were grown in Dulbecco’s minimum essential medium (DMEM) supplemented with 10% FBS (VWR Life Science, Seradigm). C3A, HepG2, SNU398, PLC/PRF/5, Hep3B2.1-7, SKHep1, SNU475, SNU387, SNU423 and SNU449 were grown in the ATCC-recommended medium. JHH4 cells were grown in Eagle’s minimum essential medium (MEM), JHH6 cells in William’s E medium and NCI-H684, SNU761, SNU886 and SNU878 in RPMI 1640 medium all supplemented with 10% FBS. For SILAC labeling, cells were grown in custom -Lys/-Arg DMEM (Caisson Labs, North Logan, UT) supplemented with 10% dialyzed FBS (10,000 Da cutoff, Sigma, St Louis, MO), Penicillin-Streptomycin-Glutamine (100x), 0.174 mM proline, 0.8 mM Lys-^13^C_6_^15^N_2_ and 0.4 mM, Arg-^13^C_6_,^15^N_4_ (Cambridge Isotope Labs, Andover, MA) for HCT-116, SH-SY5Y and U-2 OS cells or in custom -Lys/-Arg RPMI-1640 supplemented with 10% dialyzed FBS, Penicillin-Streptomycin-Glutamine (100x), 0.174 mM proline, 0.219 mM Lys-^13^C_6_^15^N_2_ and 1.14 mM Arg-^13^C_6_,^15^N_4_ for Du4475, HL-60, Jurkat and K562 cells. Cells were grown for at least 5 cell doublings in SILAC medium and harvested when reaching 90% confluency or a density of 1×10E6 cells/ml.

##### Human HCC and normal adjacent liver specimens

Primary human HCCs with paired non-tumor livers were obtained from patients undergoing liver resection at the University of Washington Medical Center (Seattle, WA, USA). All patients in this study prospectively consented to donate liver tissue for research under the Institutional Review Board protocols #1852. Once the specimens were collected under the direction of Pathology representatives, they were snap-frozen in liquid nitrogen and stored at −80°C until further processing.

### Method details

#### RNAi knockdown experiments

All lentiviral vectors encoding different shRNAs (STAT3, FYN, FZD2, AXL, NUAK1 and 2) in a pGIPZ vector were purchased from OpenBiosystems (Dharmacon, Lafayette, CO). Cell lines were transfected with shRNA constructs using Lipofectamine 2000 (Invitrogen, Carlsbad, CA) and 48 h post-transfection selected with 4 µg/ml puromycin (Invitrogen). The clones were sorted by FACS and screened for target mRNA knockdown by Western blot or qPCR analysis (see ‘Western blot analysis and antibodies’ and ‘Quantitative real-time PCR (qPCR) analysis of mRNA expression’ below). Stable cell lines were maintained in DMEM (see ‘Cell lines and cell culture conditions’) supplemented with 2 µg/ml puromycin.

#### Ectopic expression of NUAK1

The expression construct encoding for full length NUAK1 (NM_ 014840.2) in a lentiviral plasmid (pReceiver-Lv242) was purchased from Genecopoeia (Rockville, MD). Cell lines were transfected with NUAK1 plasmid construct using Lipofectamine 2000 (Invitrogen, Carlsbad, CA) and 48 h post-transfection selected in 4 µg/ml puromycin (Invitrogen). Stable cell lines were maintained in DMEM (see ‘Cell lines and tissue culture conditions’) supplemented with 2 µg/ml puromycin.

#### Western blot analysis and antibodies

Antibodies used were anti-phospho-Stat3 (Tyr705) (Cell signaling Technology, Cat # 9145), anti-E-cadherin (Cell signaling Technology, Cat # 3195), anti-Occludin (BD Transduction Laboratories, Cat # 611091), anti-Vimentin (Millipore, Cat #CS207806), anti-β-Actin (Sigma, Cat #A1978), anti-GAPDH (Santacruz Biotechnology, Cat # Sc-365062), anti-phospho-p44/42 MAPK (Erk1/2) (Thr202/Tyr204) (Cell signaling Technology, Cat #4370), anti-Axl (C89E7) (Cell signaling Technology, Cat #8661). Briefly, cells were rinsed in phosphate buffered saline (PBS) and lysed in lysis buffer (20 mM Tris-HCl, 150 mM NaCl, 1% Triton X-100 (v/v), 2 mM EDTA, pH 7.8 supplemented with 1 mM sodium orthovanadate, 1 mM phenylmethylsulfonyl fluoride (PMSF), 10 µg/ml aprotinin, and 10 µg/ml leupeptin). Protein concentrations were determined using the BCA protein assay (Pierce, Rockford, IL) and immunoblotting experiments were performed using standard procedures. For quantitative immunoblots, primary antibodies were detected with IRDye 680-labeled goat-anti-rabbit IgG or IRDye 800-labeled goat-anti-mouse IgG (LI-COR Biosciences, Lincoln, NE) at 1:5000 dilution. Bands were visualized and quantified using an Odyssey Infrared Imaging System (LI-COR Biosciences).

#### Quantitative real-time PCR (qPCR) analysis of mRNA expression

Cells were seeded in 6-well plates 24 h prior to isolation of total RNA using a RNeasy Mini Kit (QIAGEN, Santa Clara, CA). mRNA levels for EMT-related genes were determined using the validated primer sets (SA Biosciences Corporation, Frederick, MD). Briefly, 1 µg of total RNA was reverse transcribed into first strand cDNA using an RT^2^ First Strand Kit (SA Biosciences). The resulting cDNA was subjected to qPCR using human gene-specific primers for EMT-associated genes, and two housekeeping genes (GAPDH and ACTB). The qPCR reaction was performed with an initial denaturation step of 2 min at 95°C, followed by 5 s at 95°C and 30 s at 60°C for 40 cycles using an Biorad CFX384 system (Biorad, Hercules, CA). The mRNA levels of each gene were normalized relative to the mean levels of the two housekeeping genes and compared with the data obtained from unstimulated, serum-starved cells using the 2-ΔΔCt method. According to this method, the normalized level of a mRNA, X, is determined using Equation **1**:

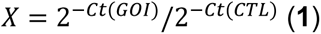

where Ct is the threshold cycle (the number of the cycle at which an increase in reporter fluorescence above a baseline signal is detected), GOI refers to the gene of interest, and CTL refers to a control housekeeping gene. This method assumes that Ct is inversely proportional to the initial concentration of mRNA and that the amount of product doubles with every cycle.

#### Kinetic wound healing assay

A wound healing assay was used to study the effect of AXL and NUAK1/2 RNAi knockdown on FOCUS cell migration and to score the cell motility of the 15 wt HCC cell lines. Briefly, cells were plated on 96-well plates (Essen Image Lock, Essen Bioscience, Ann Arbor, MI), and a wound was scratched with a wound scratcher (Essen Instruments). Wound confluence was monitored with Incucyte Live-Cell Imaging System and software (Essen Instruments). Wound closure was observed every 2 hours for 24-72 hours by comparing the mean relative wound density of at least three biological replicates in each experiment.

#### High throughput growth inhibition assay

The high throughput growth inhibition assay was performed by the QUELLOS high throughput screening facility of the University of Washington (http://depts.washington.edu/iscrm/quellos/) using 299 compounds of a Selleckchem kinase inhibitor Library (http://www.selleckchem.com/screening/kinase-inhibitor-library.html, Selleckchem, Houston, TX). Briefly, the assay was performed in a 384-well plate format in biological duplicate. Compounds were applied at 7 different concentrations ranging from 10 µM to 10 nM and cell viability was measured after 72 h of incubation using the CellTiter-Glo 2.0 assay (Promega, Madison, WI).

#### Small-molecule inhibitor and AXL/NUAK RNAi co-treatment

1800 cells/well were seeded onto white flat bottom half area 96-well plates (Greiner Bio-One, Kremsmuenster, AT) in 50 µl of growth medium and allowed to attach in an incubator for 24 h. Then the drugs in DMSO and/or DMSO vehicle controls as 12X solutions in growth medium were added to a total volume of 60 µl and 0.1% DMSO final. The cells were grown in an incubator for another 72 h. Then, 60 µl of CellTiter-Glo 2.0 (Promega, Madison, WI) reagent/well were added according to the manufacturer’s instructions and luminescence was quantified with a SpectraMax 190 plate reader (Molecular Devices, San Jose, CA). The CHEK1/2 inhibitor AZD7762 (Selleckchem, Houston, TX) and the NUAK1/2 inhibitor WZ4003 (Tocris Bioscience, Minneapolis, MN) were applied at 8 different concentrations between 10 µM and 4.6 nM (3-fold dilution steps) and Dinaciclib (Selleckchem) was applied at 8 different concentrations between 1 µM and 0.5 nM (3-fold dilution steps). Experiments were performed in four biological replicates. Growth inhibition curves were fitted using the GraphPad Prism software package (V5.0a) with a least-squares nonlinear regression model for curve fitting (One site - Fit logIC50 function).

#### Protein extraction from human HCC and normal adjacent liver specimens

Frozen tumor specimens of ca. 100 mg wet weight were ground into a fine powder using the CryoGrinder Kit from OPS Diagnostics (Lebanon, NJ). The powder was then added to ice cold mod. RIPA buffer containing phosphatase and protease inhibitors (see ‘Kinase affinity enrichment and on-bead digestion’ below), vortexed 10 times at max. speed and clarified at 21,000 rcf and 4°C for 20 min. Protein yields from specimens ranged from 5% to 10% depending on the degree of fibrosis.

#### IMAC phosphopeptide enrichment

IMAC phosphopeptide enrichment was performed according to the published protocol (in-tube batch version) with the following minor modifications (53). 20 µl of a 50% IMAC bead slurry composed of 1/3 commercial PHOS-select iron affinity gel (Sigma Aldrich, St Louis, MO), 1/3 in-house made Fe^3+^-NTA Superflow agarose and 1/3 in-house made Ga^3+^-NTA Superflow agarose were used for phosphopeptide enrichment (54). The IMAC slurry was washed three times with 10 bed volumes of 80% aq. ACN containing 0.1% TFA and phosphopeptide enrichment was performed in the same buffer.

#### Peptide and phosphopeptide desalting with StageTips

Peptides and phosphopeptides were desalted using C18 StageTips according to the published protocol with the following minor modifications for phosphopeptides (55). After activation with 50 µl methanol and 50 µl 80% aq. ACN containing 0.1% TFA the StageTips were equilibrated with 50 µl 1% aq. formic acid. Then the peptides that were reconstituted in 50 µl 1% aq. formic acid were loaded and washed with 50 µl 1% aq. formic acid. The use of 1% formic acid instead of 5% aq. ACN containing 0.1% TFA reduces the loss of highly hydrophilic phosphopeptides.

#### Preparation of optimized kinobead mixture

The seven kinobead affinity reagents used were synthesized in-house as described previously (56, 57). For optimal coverage of the human kinome an optimized mixture of the seven kinobead reagents was prepared as follows: 1 ml of reagent **1**, 0.5 ml of reagents **2**, **3** and **7**, respectively, and 0.25 ml of reagents **4**, **5** and **6**, respectively, to give 3.25 ml of the complete kinobead mixture. All reagents were a 50% slurry in 20% aq. ethanol.

#### Kinase affinity enrichment and on-bead digestion

Three micro tubes containing 35 µl of a 50% slurry of the in-house-made, optimized kinobead mixture in 20% aq. ethanol were prepared for each pulldown experiment. The beads were washed twice with 300 µl modified RIPA buffer (50 mM Tris, 150 mM NaCl, 0.25% Na-deoxycholate, 1% NP-40, 1 mM EDTA and 10 mM NaF, pH 7.8). 1 mg of protein extract in mod. RIPA buffer containing HALT protease inhibitor cocktail (100x, Thermo Fisher Scientific, Waltham, MA) and phosphatase inhibitor cocktail II and III (100x, Sigma-Aldrich, St Louis, MO) were added to the first tube. The mixture was incubated on a tube rotator for 1h at 4°C and then the beads were spun down rapidly at 2000 rpm on a benchtop centrifuge (5s). The supernatant was pipetted into the next tube with kinobeads for the second round of affinity enrichment. The procedure was repeated once more for a total of three rounds of affinity enrichment. After removal of the supernatant, the beads were rapidly washed twice with 300 µl of ice-cold mod. RIPA buffer and three times with 300 µl ice-cold tris-buffered saline (TBS, 50 mM tris, 150 mM NaCl, pH 7.8) to remove detergents. 100 µl of the denaturing buffer (20% trifluoroethanol (TFE)(58), 25 mM Tris containing 5 mM tris(2-carboxyethyl)phosphine hydrochloride (TCEP*HCl) and 10 mM chloroacetamide (CAM), pH 7.8), were added and the slurry vortexed at low speed briefly. At this stage, kinobeads from the three tubes are combined and heated at 95°C for 5 min. The mixture was diluted 2-fold with 25 mM triethylamine bicarbonate (TEAB), the pH adjusted to 8-9 by addition 1 N aq. NaOH; 5 µg LysC were added and the mixture agitated on a thermomixer at 700 rpm at 37°C for 2 h. Then 5 µg MS-grade trypsin (Thermo Fisher Scientific, Waltham, MA) were added, and the mixture agitated on a thermomixer at 700 rpm at 37°C overnight. 600 µl of 1% formic acid was added and the mixture acidified by addition of another 6 µl of formic acid to yield 1.2 ml peptide solution in total. An aliquot of 120 µl (10%) of the peptide solution was desalted using StageTips and analyzed in single nanoLC-MS/MS runs for protein quantification. The remaining peptide solution (90%) was dried under vacuum at RT on a SpeedVac. 300 µl of 70% aq. ACN + 0.1 % TFA was added to each tube, the mixture vortexed, and sonicated in a bath sonicator until dried peptide residue dissolved. In case the dried residue could not be fully resuspended, additional 0.1% aq. TFA can be added in 10 µl increments until dissolved. The solution was subjected to IMAC phosphopeptide enrichment protocol and desalted using StageTips (see ‘IMAC phosphopeptide enrichment’ and ‘Peptide and phosphopeptide desalting with StageTips’ above).

#### Kinase quantification in HCC specimens with super SILAC

Lys8/Arg10-labeled SILAC cells (HCT-116, SH-SY5Y, U-2 OS, Du4475, HL-60, Jurkat and K562) were lysed in mod. RIPA buffer and clarified as described above (‘Kinase affinity enrichment, on-bead digestion and phosphopeptide enrichment’). The heavy-labeled SILAC master mix was prepared by mixing the lysates of the seven cell lines in equal amounts. Lysate from patient tumor and NAL samples was mixed 1:1 (total amount of protein 300 µg) and subjected to kinase affinity enrichment and LC-MS/MS analysis according to the previously published protocol (57, 59).

#### nanoLC-MS/MS analyses

The LC-MS/MS analyses were as previously described (57) with minor modifications: Peptide samples were separated on a Thermo-Dionex RSLCNano UHPLC instrument (Sunnyvale, CA) using 20 cm fused silica capillary columns (100 µm ID) packed with 3 μm 120 Å reversed phase C18 beads (Dr. Maisch, Ammerbuch, DE). For whole peptide samples the LC gradient was 120 min long with 10−35% B at 300 nL/min. For phosphopeptide samples the LC gradient was 120 min long with 3−30% B at 300 nL/min. LC solvent A was 0.1% aq. acetic acid and LC solvent B was 0.1% acetic acid, 99.9% acetonitrile. MS data was collected with a Thermo Fisher Scientific Orbitrap Elite (kinobead/LC-MS experiments, global phosphoproteomics analyses) or Orbitrap Fusion Lumos Tribrid MS (global proteome analyses). Top15 selection with CID fragmentation was used for MS2 data-dependent acquisition.

#### Analysis of MS data files

Data .raw files were analyzed by MaxQuant/Andromeda (60) version 1.5.2.8 using protein, peptide and site FDRs of 0.01 and a score minimum of 40 for modified peptides, 0 for unmodified peptides; delta score minimum of 17 for modified peptides, 0 for unmodified peptides. MS/MS spectra were searched against the UniProt human database (updated July 22nd, 2015). MaxQuant search parameters: Variable modifications included Oxidation (M) and Phospho (S/T/Y). Carbamidomethyl (C) was a fixed modification. Max. missed cleavages was 2, enzyme was Trypsin/P and max. charge was 7. The MaxQuant “match between runs” feature was enabled. The initial search tolerance for FTMS scans was 20 ppm and 0.5 Da for ITMS MS/MS scans.

#### MaxQuant output data processing

MaxQuant output files were processed, statistically analyzed and clustered using the Perseus software package v1.5.6.0 (61). Human gene ontology (GO) terms (GOBP, GOCC and GOMF) were loaded from the ‘Perseus Annotations’ file downloaded on 01.08.2017. Expression columns (protein and phosphopeptide intensities) were log2 transformed and normalized by subtracting the median log2 expression value from each expression value of the corresponding data column. Potential contaminants, reverse hits and proteins only identified by site were removed. Reproducibility between LC-MS/MS experiments were analyzed by column correlation (Pearson’s r) and replicates with a variation of r > 0.25 compared to the mean r-values of all replicates of the same experiment (cell line or knockdown experiment) were considered outliers and excluded from the analyses. Data imputation was performed using a modeled distribution of MS intensity values downshifted by 1.8 and having a width of 0.2. For statistical testing of significant differences in expression, a two-sample Student’s T-test with Benjamini-Hochberg correction for multiple hypothesis testing was applied (FDR = 0.05). For statistical testing of the 17 HCC cell line data (EMT state-association) all biological replicates were used. For MS protein intensities this was n = 42 (epithelial cells) and n = 60 (mesenchymal cells). For MS phosphopeptide intensities this was n = 56 (epithelial cells) and n = 91 (mesenchymal cells).

#### Co-expression analysis of LIHC mRNA data

mRNA intensity values for the 101 drug EMT state-associated kinases identified by kinobead/LC-MS analysis of the 17 HCC cell line panel (**Table S2**) and the 50 EMT and stemness markers (**Table S1**) were extracted from the TCGA-LIHC dataset (13) mRNA intensities were log2 transformed and pairwise correlation analysis was performed separately for mesenchymal and epithelial mRNAs across all tissue samples in the dataset (n = 424) using Pearson’s r. The resulting symmetrical matrices of r-values were subjected to hierarchical clustering using R (gplots::heatmap.2) to identify groups of co-expressed mRNAs (see Figure 4D and 4E).

#### Pharmacoproteomic AUC–MS intensity correlation

Mean LFQ-MS intensities values for each of the 17 cell lines were calculated using the imputed MS intensity data from **Table S2**. All proteomics features (n = 13935) were correlated with each compound’s AUC value across the cell line panel (n = 17) using the Pearson’s correlation coefficient resulting in a 13935×299 matrix of r-values (see **Table S4**). To rank proteomics feature the resulting r-values for each kinase inhibitor were sorted from low to high where negative r-values correspond to drug sensitivity (low AUC - high MS intensity) and positive r-values correspond to drug resistance (high AUC–high MS intensity).

#### Kinome-GSEA analysis with Reactome pathways

To obtain signaling pathways for GSEA analyses, we used the mapped identifier files ‘NCBI2Reactome_All_Levels.txt’ and ‘ReactomePathwaysRelation.txt’ from Reactome.org (downloaded 22th Oct., 2018). Pathways from the highest hierarchical levels were removed to exclude non-specific pathways. Subsequently, a regular expression match for patterns containing “kinase”, “signal”, “cell cycle”, “migration”, “cancer”, “dna repair”, “mitos” and “mitot” was used to extract 327 cancer relevant pathways (**Table S3** ‘Reactome_Pathways’). Member genes of each pathway were mapped to unique identifiers in **Table S3** to allow addition of phosphopeptide data. The Pearson correlation coefficients r from correlating drug response with MS intensities of kinome features (see ‘Pharmacoproteomic AUC – MS intensity correlation analyses’ above) were –(x) transformed for GSEA analyses, as the GSEA algorithm ranks features by their correlation with drug response. We used the Bioconductor package, fgsea (62) (https://doi.org/10.1101/060012), with parameters: minSize = 10, maxSize = 500, gseaParam = 2 and nperm = 10000 to compute p-values and enrichment scores, including corrections for multiple hypothesis testing (BH FDR = 0.1).

#### Classification of KI drugs by correlation-clustering of kinome-GSEA wNES scores

For hierarchical clustering of compounds based on Reactome pathway analysis (Figure S4D), FDR-weighted Reactome pathway normalized enrichment scores (wNES = NES*(1-FDR)) were extracted for each compound (N = 299, **Table S3** worksheet ‘299_Compound_NES_FDR’). Pairwise Pearson correlation coefficients (r) for wNES values for all compounds were calculated resulting in a 299X299 matrix of r-values. Clustering of the Pearson’s correlation coefficients using R (gplots::heatmap.2) identified 11 major groups (Figure 4D and **Table S3** tab ‘Reactome_Pathway_Clusters’). For these 11 major groups we then calculated mean wNES values for 34 cancer-relevant, non-redundant Reactome pathways representative of the larger panel of Reactome pathways (Figure 3A). These 34 pathways were selected for having the largest difference in mean wNES across all 11 KI drug clusters among Reactome terms in the same overarching pathway theme. The result is a 11X34 matrix of mean wNES values (Figure 3A) that was subjected to hierarchical clustering using R (gplots::heatmap.2).

#### Kinome-GSEA in clinical HCC specimens and correlation of Pathway signatures

We calculated the difference of the mean imputed MS intensity values for the protein and phosphopeptide expression kinobead LFQ-MS data (**Table S5**) tumor vs. normal adjacent liver and ranked all proteomics features according to this difference of the mean (positive values was enriched in tumor, negative values was enriched in). We then applied our kinome-GSEA analysis and calculated FDR-weighted Reactome pathway enrichment scores (wNES values, see ‘Kinome-GSEA analysis with Reactome pathways’). We then calculated Pearson correlation coefficients, correlating the wNES values of all 299 KI drugs with the wNES values of the four HCC tumor/NAL pairs. High r-value then indicate enrichment of a KI pathway signature in tumors, and negative values their enrichment in NAL.

#### Comparison of mRNA-GSEA and kinome-GSEA performance

First, we assumed that enrichment of a target Reactome pathway of a KI drug (that is a Reactome pathway that contains the official gene name of a target kinase) by our mRNA-GSEA and kinome-GSEA analyses is an indicator of successful prediction of KI drug response. KI drugs were binned into 39 primary kinase target groups (**Table S3**) according to their known molecular targets as is indicated by the commercial provider (Selleckchem) with a cutoff of EC_50_ < 500 nM for a given target kinase. We then extracted FDR-weighted NES values (wNES) from **Table S3** for compounds in each of the primary target groups with ≥5 members (N = 27, see Figure S3). We calculated the mean wNES across all Reactome pathways that contain the gene name of the primary kinase target of that group (target Reactome pathways). This was done separately for mRNA-GSEA and kinome-GSEA and the results plotted in a dot-line plot (Figure S3). Positive mean wNES values for a compound indicate enrichment of target Reactome pathways and thus prediction of the correct target pathway. For selected examples of important kinase inhibitor groups identified by kinome-GSEA pathway clustering the mean wNES values were plotted in a box plot for comparison (Figure 2A). Where the mean wNES is more positive, either mRNA-GSEA or kinome-GSEA, we assumed the corresponding method more predictive of KI drug response.

#### Functional phosphosites and kinase-substrate relationships

To determine the biological function of a phosphorylation site and the kinase-substrate relationship of a given phosphorylation site the PhosphoSite Plus datasets ‘Regulatory_Sites’ and ‘Kinase_Substrate_Dataset’ were searched against the 15 amino acid sequence windows centered on the corresponding phosphosite. Human, mouse and rat phosphorylation sites were all considered to assess the biological and biochemical consequences of phosphorylation. The datasets were downloaded from the PhosphoSite Plus webpage on the 13^th^ of March, 2017 (https://www.phosphosite.org/)(27).

#### Kinome dendrograms

Kinome dendrograms were prepared using the KinMap web application (http://kinhub.org/kinmap/)(63)..

#### Identification of protein kinase interactors

Protein kinase interactors were determined using the BioGRID database only considering protein-protein interactions for which two independent lines of evidence exist (26). To that end, the ‘BIOGRID-MV-Physical-3.5.165.tab2’ file was downloaded on October 6^th^, 2018 and mined for protein kinase interactions through matching against the gene name in the MaxQuant output files.

#### Construction of interaction network graphs

Protein-protein interaction network graphs were plotted with the STRING web application (v10.0, https://string-db.org/)(51). Solid edges shown in network figures represent the ‘confidence’ in the existence of a physical interaction

#### Quantitation and statistical analysis

Differences between sample populations were quantified with a two-tailed two sample Student’s T-test. For testing of proteomics data or where indicated BH correction for multiple hypothesis testing (FDR = 0.05) was applied.

#### Data and software availability

MS raw files and MaxQuant/Andromeda output files were deposited in the MassIVE repository under the dataset ID: MSV000083236. Drug response–kinase pathway interaction data can be viewed interactively via our Shiny web application (*25*); this resource allows users to view the association of signaling pathways, kinome features, and kinase inhibitors across the complete dataset.

### Supplementary Table Titles and Legends

**Table S1. HCC Cell EMT Marker, Kinase mRNA and Drug Panel; Supplement to Figure 1-4.** Log2 mRNA intensity of 50 EMT and stem cell markers as well as 509 protein kinases in the 28 CCLE HCC cell lines. Names and primary targets of the 299 kinase inhibitor panel.

**Table S2. Kinome Profiling 17 HCC Cell Lines; Supplement to Figure 1-3.** MaxQuant protein groups and phospho(S/T/Y) output data obtained from kinome profiling of 17 HCC cell lines. Missing values were replaced by data imputation (see ‘Materials and Methods’).

**Table S3. Kinase Inhibitor HTS Results and kinome-GSEA Analysis; Supplement to Figure 1-3.** Results of the HTS of 299 kinase inhibitors in 17 HCC cell lines. Summary table of AUC values. Summary of Reactome pathways used in the kinome-GSEA analysis. Compounds and cumulative NES for clusters from **Figure S2D**. FDR and NES values for all 299 compounds and 275 Reactome pathways.

**Table S4. Kinome Proteomic Feature-AUC Correlation Results; Supplement to Figure 1-3.** Results from Pearson correlation between mean MS intensity values of all quantified proteomics features (n = 13935) and kinase inhibitor AUCs (n = 299) across the 17 HCC cell line panel.

**Table S5. Kinome Profiling of Human HCC Specimens; Supplement to Figure 4.** Results of protein kinase quantification by super SILAC kinobead/LC-MS profiling in 9 paired (tumor/NAL) human HCC specimens. MaxQuant protein groups and phospho(S/T/Y) output data from LFQ kinobead/LC-MS profiling of 4 paired (tumor/NAL) human HCC specimens. Kinome-GSEA enrichment in clinical HCC specimens.

**Table S6. Kinome Profiling FOCUS RNAi Models; Supplement to Figure 5 and 6.** MaxQuant protein groups and phospho(S/T/Y) output data based on kinome profiling of the FOCUS cell AXL, FZD2 and NUAK1/2 RNAi models.

